# Microbiome induced complement synthesized in the gut protects against enteric infections

**DOI:** 10.1101/2023.02.02.523770

**Authors:** Meng Wu, Wen Zheng, Xinyang Song, Bin Bao, Yuanyou Wang, Deepshika Ramanan, Daping Yang, Rui Liu, John C. Macbeth, Elyza A. Do, Warrison A Andrade, Tiandi Yang, Hyoung-Soo Cho, Francesca S. Gazzaniga, Marit Ilves, Daniela Coronado, Charlotte Thompson, Saiyu Hang, Isaac M. Chiu, Jeffrey R. Moffitt, Ansel Hsiao, John J. Mekalanos, Christophe Benoist, Dennis L. Kasper

## Abstract

Canonically, complement is a serum-based host defense system that protects against systemic microbial invasion. Little is known about the production and function of complement components on mucosal surfaces. Here we show gut complement component 3 (C3), central to complement function, is regulated by the composition of the microbiota in healthy humans and mice, leading to host-specific gut C3 levels. Stromal cells in intestinal lymphoid follicles (LFs) are the predominant source of intestinal C3. During enteric infection with *Citrobacter rodentium* or enterohemorrhagic *Escherichia coli,* luminal C3 levels increase significantly and are required for protection. *C. rodentium* is remarkably more invasive to the gut epithelium of C3-deficient mice than of wild-type mice. In the gut, C3-mediated phagocytosis of *C. rodentium* functions to clear pathogens. Our study reveals that variations in gut microbiota determine individuals’ intestinal mucosal C3 levels, dominantly produced by LF stromal cells, which directly correlate with protection against enteric infection.

**Highlights:**

1. Gut complement component 3 (C3) is induced by the microbiome in healthy humans and mice at a microbiota-specific level.
2. Gut stromal cells located in intestinal lymphoid follicles are a major source of luminal C3
3. During enteric infections with *Citrobacter rodentium* or enterohemorrhagic *Escherichia coli,* gut luminal C3 levels increase and are required for protection.
4. *C. rodentium* is significantly more invasive of the gut epithelium in C3-deficient mice when compared to WT mice.
5. In the gut, C3-mediated opsonophagocytosis of *C. rodentium* functions to clear pathogens.

## Introduction

Infectious diarrheal disease is the second leading cause of childhood death worldwide, and death rates are especially high in low- and middle-income countries.^1^ On average, diarrhea kills 2,195 children every day—more than AIDS, malaria, and measles combined.^2^ Out of the 1.7 billion cases of childhood diarrheal disease that occur globally, approximately 500,000 cases result in death.^3^ Furthermore, diarrhea also has a detrimental impact on growth and cognitive development due to the disruption of both intestinal absorptive and barrier functions. These problems bestow great urgency to the development of new and improved approaches to combat enteric infections.^4^

The complement system comprises more than 50 secreted proteins and membrane receptors that function in a highly coordinated fashion to form one of the first lines of innate immune defense against invasion of host tissues by microbes. Complement is classically considered a chiefly hepatocyte-derived and serum-resident system, acting in the blood and interstitial fluids.^5^ Complement can be activated through the classical, lectin, and alternative pathways, which converge at complement component 3 (C3), the central element of the complement cascade. Deposition of a fragment of C3 (C3b) on the surface of bacterial cells is essential for direct lysis of many gram-negative bacteria by driving formation of the membrane-attack complex (MAC) or by promoting phagocytosis and killing by neutrophils and macrophages.^6^ Surprisingly little is known about the function, if any, of the complement system on the apical side of epithelial barriers (e.g., in mucosal secretions of the respiratory, urinary, and gastrointestinal tracts), where some of the earliest encounters with potentially invading bacterial pathogens take place.

During the COVID-19 pandemic, studies found that the complement system is one of the most highly induced pathways in lung epithelial cells following SARS-CoV-2 infection.^7^ Complement cascade genes are also enriched among differentially upregulated genes in rectal biopsies from patients with inflammatory bowel disease (IBD) relative to healthy controls.^8^ However, the scale and function of the complement system in the gastrointestinal tract is unclear, despite the gut representing the primary reservoir of host-associated microbes. The gut microbiota (the community of bacteria, archaea, fungi, protists, and viruses that inhabit the intestine) plays a critical role in many aspects of human health and may modulate susceptibility to a wide spectrum of diseases such as IBD, obesity, diabetes, neurologic disease, malnutrition, and diarrheal infections.^8–12^ However, the role of the intestinal microbiota in modulating complement activity in the gut mucosa is poorly understood.

In this study, we show that complement C3 is synthesized in the gut by three key types of cells and secreted into the intestinal lumen, and that the luminal C3 level is determined by the composition of the host’s gut microbiota both in humans and mice. We demonstrate that secreted luminal C3 is critical for the host to fight against infectious diarrhea due to enteric pathogens such as enterohemorrhagic *Escherichia coli* (EHEC) and *Citrobacter rodentium*, and that low gut C3 levels serve as a risk factor for enteric infection. We identified stromal cells as the primary source of gut complement C3 during homeostasis, and that these cellular populations potently upregulated C3 expression during infection. This microbiota-dependent complement C3 production in the gut underpins a relationship between specific gut microbes and the pathogenesis of infectious diarrhea in children and adults. Further understanding of these interactions may provide crucial insight into the pathogenesis, treatment, and prevention of infectious diarrhea, one of the most significant and serious health problems in children.

## Results

### C3 is produced locally within the intestine in a microbiota-dependent manner

To investigate the role of the commensal gut microbiota in host complement production, we measured C3 in feces and serum of age- and gender-matched wild-type (WT) germ-free (GF) and specific-pathogen-free (SPF) C57BL/6 mice by ELISA (Figure 1A). We found that SPF mice have significantly higher levels of intestinal C3 than GF animals (Figure 1B) while the C3 levels in the serum of GF mice were comparable to that of SPF control mice (Figure 1C). This result suggested that there are microbiota-dependent C3 differences in the gut lumen, and these differences were not due to systemic changes in serum complement levels. Gut C3 production and/or secretion are different in GF and SPF mice perhaps because of a process occurring within the intestinal mucosa.

**Figure 1:**
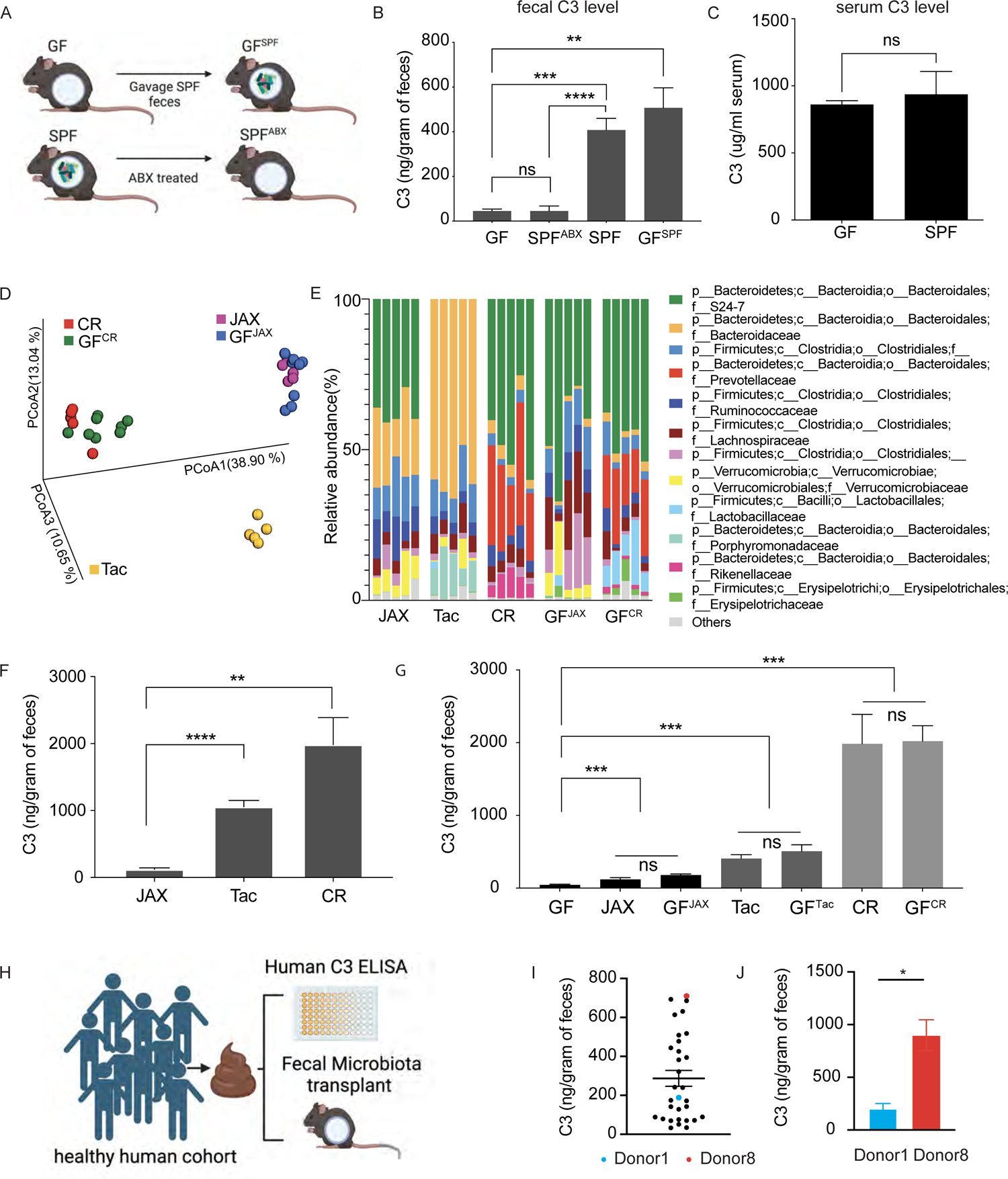
Gut complement factor C3 is regulated by the host microbiota. (A) Mouse models used to evaluate fecal C3 levels in (B). GF: C57BL/6 germ-free mice; SPF: specific-pathogen free C57BL/6 mice harbor a normal murine microbiota; SPF^ABX^: SPF mice treated with antibiotics for two weeks; GF^SPF^: previously germ-free mice colonized with SPF microbiota. (B) Fecal C3 levels in GF, SPF^ABX^, SPF, and GF^SPF^. (C) C3 levels in serum of GF and SPF mice. (D) Principal Coordinate Analysis (PCoA) of unweighted UniFrac distance measurements based on the 16S rRNA gene sequencing data of the composition of bacterial communities in fecal samples collected from C57BL/6 mice from three commercial vendors: Jackson (JAX); Taconic (Tac); and Charles River (CR) or GF mice colonized with JAX (GF^JAX^) or CR microbiota (GF^CR^) ((q-values=0.001, PERMANOVA). (E) Family level abundance analysis of fecal microbiota from the mice in Figure 1D. (F) Fecal C3 levels in C57BL/6 mice from three commercial vendors: Jackson (JAX); Taconic (Tac); and Charles River (CR). (G) Fecal C3 levels of gnotobiotic mice colonized for two weeks with fecal samples from the indicated commercial sources. (H) Experimental scheme to evaluate fecal C3 levels in a healthy human cohort and their relationship to the gut microbiota. (I) Fecal C3 levels in 31 healthy adult human donors. (J) Fecal C3 levels in gnotobiotic mice colonized with fecal microbiota from human donor 1 or donor 8. ns: not-significant,*p<0.05, **p<0.01, *** p<0.001, **** p<0.0001, unpaired student’s t-test.

To further test whether the production of C3 in the lumen could be triggered by exposure to microbes post-weaning or is programmed during early development, we colonized 4-week-old GF mice with SPF microbiota (Figure 1A). Two weeks after introducing an SPF mouse fecal slurry into GF mice (generating GF^SPF^ mice), there were comparable levels of C3 in fecal specimens from the GF^SPF^ mice as in fecal samples of SPF animals. We examined whether removing members of the gut microbiota reduced fecal C3 levels to those observed in GF mice. After two weeks of broad-spectrum antibiotic treatment in the drinking water, antibiotic-treated SPF mice (SPF^ABX^) had no detectable live bacteria in their feces by culturing, and fecal C3 levels were like those of GF mice (Figure 1B).

We investigated whether luminal C3 levels were affected by the composition of the microbiota or simply reflected a generic response to colonization with any microbe. We measured C3 levels in WT 8-10-week-old C57BL/6 mice from three commercial breeders: Jackson Laboratory (JAX), Taconic (Tac), and Charles River (CR). We found that inbred animals of the same strain from each vendor harbored a distinct microbiota (Figure 1D, E; Figure S1A-C) and had significantly different concentrations of fecal C3 (Figure 1F). We then transplanted the gut microbiota from each of these three mouse vendors into GF mice and observed that colonization and C3 levels recapitulated donor signatures (Figure 1G). Oral gavage of FITC-dextran in CR mice showed similar barrier permeability compared to JAX mice (Figure S1D). Blind histology check scored both JAX and CR mice at zero with no histologic sign of disease (Figure S1E). These data suggest that the composition of the microbiota regulates levels of gut C3 production and secretion.

We subsequently investigated if our findings were germane to humans. We measured fecal C3 levels from a cohort of healthy human donors without chronic gut inflammation^12^ and found substantial individual variation in C3 levels in these fecal samples (range of 33–710 ng/g of feces) (Figure 1H, I). We transplanted the fecal microbial communities from either a high C3-producing (donor 8) or a low C3-producing (donor 1) human donor into GF mice and measured C3 levels in mouse fecal specimens two weeks after colonization (Figure 1H). Importantly, gnotobiotic mice colonized with donor 1 fecal slurry had low fecal C3 levels while mice colonized with donor 8 microbiota had high fecal C3 levels (Figure 1J). These findings suggest that microbiota from different humans can induce different levels of fecal C3 in the mouse gut and that specific microbes are a factor modulating C3 levels in the gut.

### *Prevotella spp.* are one of the drivers of high-C3 production in the gut

To identify which specific bacterial species contribute to the induction of high C3 levels in SPF animals, we examined the C57BL/6 mouse microbiota of JAX and CR as representative of low-C3 and high-C3 communities, respectively. We co-housed 4-week-old JAX and CR mice for 2 weeks and found that the co-housed JAX mice had increased fecal C3 levels to that of CR mice, while the co-housed CR mice retained their original baseline C3 production (Figure 2A). 16S rRNA gene sequencing of fecal samples showed that co-housed JAX mice acquired a microbiota very similar to the CR microbiota (Figure 2B, Figure S2A-D), which suggested the presence of CR-specific strains that could induce high C3 production. To identify possible candidate strains, we employed a method we previously established known as “triangulation of microbe-phenotype relationships.”^13^ In brief, we co-housed different groups of 4-week-old JAX and CR mice for one or three days to allow for microbiota transfer between mice from these two vendors. Two weeks after separation of the co-housed animals, we measured fecal C3 levels. We found that co-housing JAX mice with CR mice for either one or three days resulted in C3 levels similar to those seen in mice gavaged with CR feces (Figure 2C). To determine microbiota composition following co-housing, fecal samples were collected from these mice and subjected to 16S rRNA gene sequencing. Overall, co-housing JAX and CR mice for one day resulted in CR-like signatures in recipient JAX mice (JAX^coh_1d^) both in terms of the presence of bacterial taxa (*α*-diversity) as well as the relative abundance of taxa (*β*-diversity) (Figure 2B and Figure S2A-D). Differential abundance analysis of the microbiota composition of JAX and JAX^coh_1d^ mice identified *Prevotella spp.*, *Rikenellaceae spp*. and *Lactobacillus salivarius* as significantly more abundant in the microbiota of JAX^coh_1d^ compared with JAX mice (Figure 2D, E, S2E, F, Table S1). This suggested that these bacteria could be associated with high fecal C3 levels and be potential high-C3 inducers.

**Figure 2:**
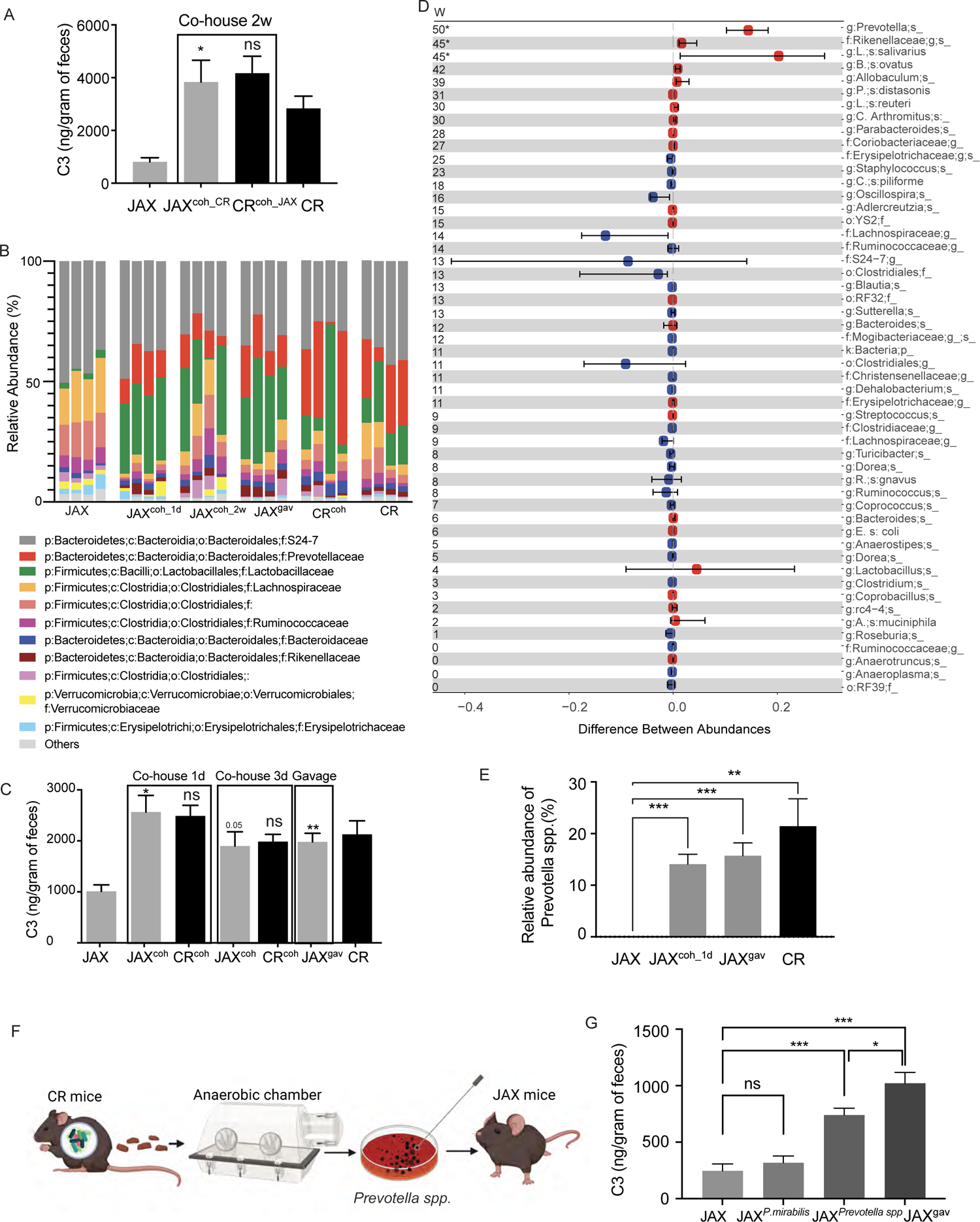
*Prevotella spp.* induce production of high luminal C3 levels in a complex host microbiota. (A) Fecal C3 levels of JAX or CR mice by themselves or co-housed for two weeks with mice from the other vendor (JAX^coh_CR^: JAX mice co-housed with CR mice, CR^coh_JAX^: CR mice co-housed with JAX mice). Statistical analyses were performed comparing treated mice to their untreated control. (B) Fecal C3 levels in JAX or CR mice co-housed for either one or three days, also in JAX mice gavaged with CR microbiota (JAX^gav^). Statistical analyses were performed comparing treated mice to their untreated control. (C) Family level abundance analysis of fecal microbiota from the mice in (A) and (B). (D) Microbiome alterations at the species level in JAX mice cohoused with CR mice for one day (JAX^coh_1d^) compared to JAX controls. Colored dots indicate the bacterial abundances differences; lines indicate the 95% confidence interval (CI) calculated using the bootstrap method. Bacterial species were identified by 16S rRNA gene sequencing and analyzed using the Wilcoxon rank-sum test. Red: Increase; Blue: Decrease. Significance was determined by W-value calculated using Analysis of Composition of Microbiomes (ANCOM) and * indicated the species with significant alterations. (E) Relative abundance of *Prevotella spp.* in JAX or CR mice, as well as in JAX mice cohoused with CR mice for one day (JAX^coh_1d^), or in JAX mice gavaged with CR microbiota (JAX^gav^). (F) Overall experimental scheme for the isolation and testing of *Prevotella spp.* Fecal samples were collected from CR mice, and resulting strains were inoculated into JAX mice to test for C3 induction in an already complex microbiota. (G) Fecal C3 levels in JAX mice compared to JAX mice colonized with either *Proteus mirabilis*, *Prevotella spp.,* or CR microbiota. ns: not-significant,*p<0.05, **p<0.01, *** p<0.001, **** p<0.0001, unpaired student’s t-test.

To identify commensal strains that could also correlate with C3 production in humans, we compared C3 levels of individual human fecal samples with commensal taxa identified by 16S rRNA gene sequencing. Due to the extensive individual variation of bacterial species, we performed a genus-level analysis. We found that the genus *Prevotella* also positively correlated with fecal C3 levels across the cohort (Spearman r=0.5487, p-value=0.029), while the genus *Actinomyces* was negatively correlated with C3 levels (Spearman r=-0.6991, p-value=0.0025) (Table S2).

Given the positive correlation between the abundance of genus *Prevotella* and high fecal C3 levels in both humans and mice, we sought to further examine its role in C3 production. We first isolated a *Prevotella spp.* (strain CR1) from CR mouse feces using selective growth conditions^14^ and colonized (by gavage) 4-week-old SPF JAX mice with CR1 (Figure 2F). We used *Proteus mirabilis (*strain CR2) isolated from CR mice as a control because it was only present in the CR microbiota; the presence of CR2 showed no significant positive correlation with C3 induction in our analysis. We found that in contrast to mice colonized with *P. mirabilis* CR2, JAX mice colonized with *Prevotella* strain CR1 showed a significant increase of fecal C3 production compared to untreated SPF JAX control mice (Figure 2G). Nevertheless, colonization with *Prevotella* CR1 did not entirely recapitulate the effect of a CR-microbiota gavage, suggesting that there are additional bacterial strains in CR mice which could induce high C3 production (Figure 2G). These data are consistent with our finding that multiple bacterial strains were identified as significantly positively associated with high fecal C3 levels in JAX^coh_1d^ mice (Figure 2D). Together, these data demonstrate that *Prevotella strain* CR1 from the CR microbiota can induce high levels of luminal C3 production in a complex microbiota environment.

### Stromal cells are the major intestinal cell population transcribing complement protein C3 during homeostasis

Although complement proteins in serum are primarily produced and secreted by the liver^15–17^, recent studies have also shown extrahepatic C3 production by various immune and non-immune cell types.^18–21^ While intestinal epithelial cells (IECs) have been suggested to produce C3 under experimental and inflammatory conditions^22–24^, the cellular source(s) for intestinal C3 at homeostasis remain unclear.

To identify which intestinal cell types express C3 under homeostasis, we used single-cell RNASeq (scRNASeq) to perform a transcriptional survey of C3 expression across intestinal populations. We used a C3-tdTomato reporter mouse line (C3^IRES-tdTomato^ C57BL/6)^25^ to isolate all colonic C3-expressing cells. Flow-cytometry analysis on colonic cells from the reporter mice found the subepithelial compartment, which includes both the lamina propria and the muscularis of the colon, as the dominant source of C3-expressing cells, compared to the epithelium (Figure 3A, Figure S3A). Therefore, we focused on the subepithelial compartment for our scRNASeq analysis. We isolated C3-expressing cells (tdTomato-positive) from the subepithelial compartment of both male and female C3^IRES-tdTomato^ C57BL/6 mice and performed scRNASeq on these cells (Figure 3B). Uniform manifold approximation and projection (UMAP) analysis revealed three major cell populations clustered by gene expression profile (Figure 3C). Using established genetic markers, we identified the three major C3-expressing populations as: 1) stromal cells (clusters 0, positive for the fibroblast cell marker podoplanin (*pdpn*), representing ∼79% of all C3-expressing colonic cells), 2) myeloid cells (clusters 1, expressing the myeloid cell marker lysozyme 2 (*lyz2*), representing ∼12% of all C3-expressing colonic cells), and 3) epithelial cells (clusters 2, highly expressing the epithelial cell marker EpCAM (*epcam*), representing 9% of all C3-expressing colonic cells) (Figure 3D).

**Figure 3:**
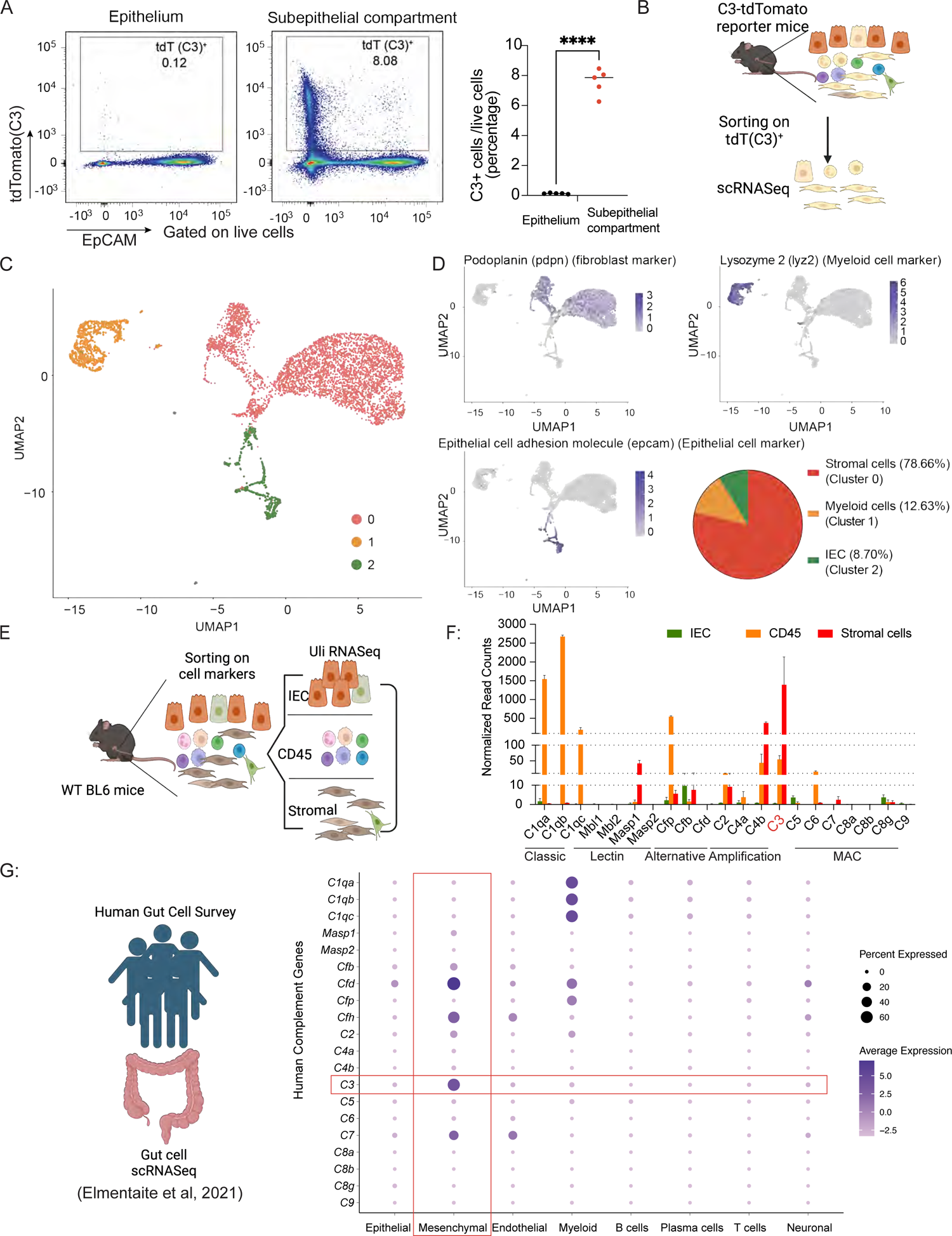
Stromal cells are the predominant cell population expressing C3 during homeostasis. (A) Representative plots of the live colonic epithelial and subepithelial compartment of C3^IRES-tdTomato^ reporter mice showing the frequency of C3-tdTomato^+^ cells and the statistics. (B) Overall experimental scheme using C3^IRES-tdTomato^ reporter mice to identify cellular sources for C3 in the colon. (C) UMAP shows scRNASeq profiles of tdTomato positive mouse colonic cells (dots) from C3^IRES-tdTomato^ reporter mice, clustered into three cell types as indicated by the colors. (D) UMAP with cells organized by cell type and colored by Pdpn, Lyz2, and Epcam expression levels. (E) Overall experimental scheme using WT C57BL/6 mice to verify cellular sources for C3 in the colon. (F) Complement system gene expression levels in WT C57BL/6 mice of the three indicated populations by Ultra Low input (Uli-RNASeq). (G) Dot plot showing average expression levels (color) of genes in the human complement system (rows) in several colonic cell populations (columns) from human adult colon samples; dot size represents the percent of cells within each parent population (column) that express these genes (row). **** p<0.0001, unpaired student’s t-test.

We then confirmed our observations by flow cytometric analysis of C3-expressing cells (tdTomato positive) from the subepithelial compartment of C3^IRES-tdTomato^ mice (Figure S3B). We confirmed that stromal cells (EpCAM^-^CD45^-^CD31^-^PDPN^+^ cells) formed the major C3-transcribing population in the subepithelial compartment under homeostatic conditions, representing 80% of the C3-expressing population. As expected from the scRNASeq data, CD45^+^ cells (∼15%) and intestinal epithelial cells (IECs) (∼5%) were the populations with the second and third highest C3 expression, respectively (Figure S3C).

To validate our reporter mouse results, we examined the indigenous C3 transcriptional levels of these three colonic cell populations using WT C57BL/6 mice. We used fluorescence-activated cell sorting (FACS) to isolate stromal (EpCAM^-^CD45^-^CD31^-^PDPN^+^ cells), CD45^+^, and IEC (EpCAM^+^) populations from the subepithelial compartment of the colon of WT C57BL/6 mice (Figure 3E). Using bulk RNA-seq analysis, we found C3 transcriptional levels in stromal cells were significantly higher than in IECs and CD45^+^ cells (Figure 3F). These results were consistent with our scRNASeq and flow-cytometry findings using C3^IRES-tdTomato^ reporter mice. Taken together, these data suggested that under homeostasis, intestinal C3 is largely expressed by stromal cells located in the subepithelial compartment of the intestine.

To identify human intestinal cellular sources for C3, we analyzed scRNASeq data from the Human Gut Cell Survey^26^, which included 79,929 intestinal cells from five anatomical sites in healthy adult human donors. B-, T-, endothelial, epithelial, mesenchymal, myeloid, neuronal, and plasma cells were identified based on their unique gene expression profiles. After retrieving the expression profile of C3 from the 79,929 intestinal cells in healthy adult human donors, we identified mesenchymal cells as the highest C3-transcribing population. Dot plot of C3 expression across cell types showed that mesenchymal cells have the largest percentage of cells expressing C3, as well as the highest average expression levels (Figure 3G). Our observations in mice are consistent with the findings using human datasets.

### Complement system gene expression profile in the intestine during homeostasis

Since complement proteins in serum usually function together in groups acting in sequence, we analyzed the intestinal transcriptional expression of complement proteins other than C3 in both mice and humans. We evaluated several major complement components at the transcriptional level using RNASeq data from the three sorted WT populations described above. Based on normalized read counts for respective genes within each population, we defined transcript expression levels as low (<10/million reads), medium (10-100/million reads), or high (>100/million reads) (Figure 3F). The analysis showed that the classical-pathway component C1q (including C1qa, C1qb, and C1qc) is highly expressed in CD45^+^ cells, consistent with a recent report that macrophages form the predominant source of C1q in the murine intestine.^27^ Interestingly, mannose-binding lectin (MBL) pathway proteins showed very low expression levels in all three populations with the exception of Masp1 (mannose-associated serine protease 1), which was expressed at a medium level in stromal cells. The alternative pathway component Cfp was highly transcriptionally expressed in CD45^+^ cells and expressed at low levels in epithelial and stromal cells. C2 and C4a exhibited very low transcriptional expression in all three cell types, while C4b was highly expressed in stromal cells and moderately expressed in CD45^+^ cells. The genes for proteins downstream of C3, including complement membrane attack complex (MAC) components C5, C6, C7, C8 and C9, showed very low or undetectable levels of transcriptional expression (Figure 3F). Overall, this analysis suggested that in the murine colon, C1q, Cfp and C4b are the only other major complement components (other than C3) with high transcriptional expression in a homeostatic environment.

Similarly to murine intestinal cells, human intestinal samples analyzed by scRNASeq also possessed high expression of classical-pathway components C1qa, C1qb, and C1qc in the CD45^+^ myeloid compartment. Human MBL pathway expression also remained equivalent to that of mice. Interestingly, alternative pathway components Cfd and Cfh were highly expressed in mesenchymal cells, and Cfp and Cfd were highly expressed in myeloid cells. While MAC member C7 was detected in mesenchymal and endothelial cells, none of the other MAC components (e.g., C5, C6, C8 or C9) were observed at detectable levels (Figure 3G). Based on these results, we conclude that mice and humans share similar expression of C3 and other main components of the gut complement system.

### C3-expressing stromal cells are primarily located in colonic lymphoid follicles and produce C3 upon bacterial stimulation

To locate C3-expressing cells in the colon, we performed RNAscope analysis using C3-specific anti-sense probes on formalin-fixed, paraffin-embedded (FFPE) sections from the colon of WT C57BL/6 mice and found that the majority of C3-expressing cells were located in the isolated lymphoid follicles (ILFs) (Figure 4A). ILFs are tertiary lymphoid organs which consist of collections of B-, T-, myeloid, and stromal cells, and their development has been suggested to be linked to microbial exposure.^28^ Morphologically, the majority of C3-expressing cells in ILFs are stromal cells. We confirmed this using immunofluorescent antibodies against tdTomato (FigureS4A-C) and PDPN on frozen colon cryo-sections from C3^IRES-tdTomato^ BL6 mice. Our staining showed co-localization of C3 and PDPN transcripts (Figure 4B) and confirmed that stromal cells in ILFs are the major cell population expressing C3.

**Figure 4:**
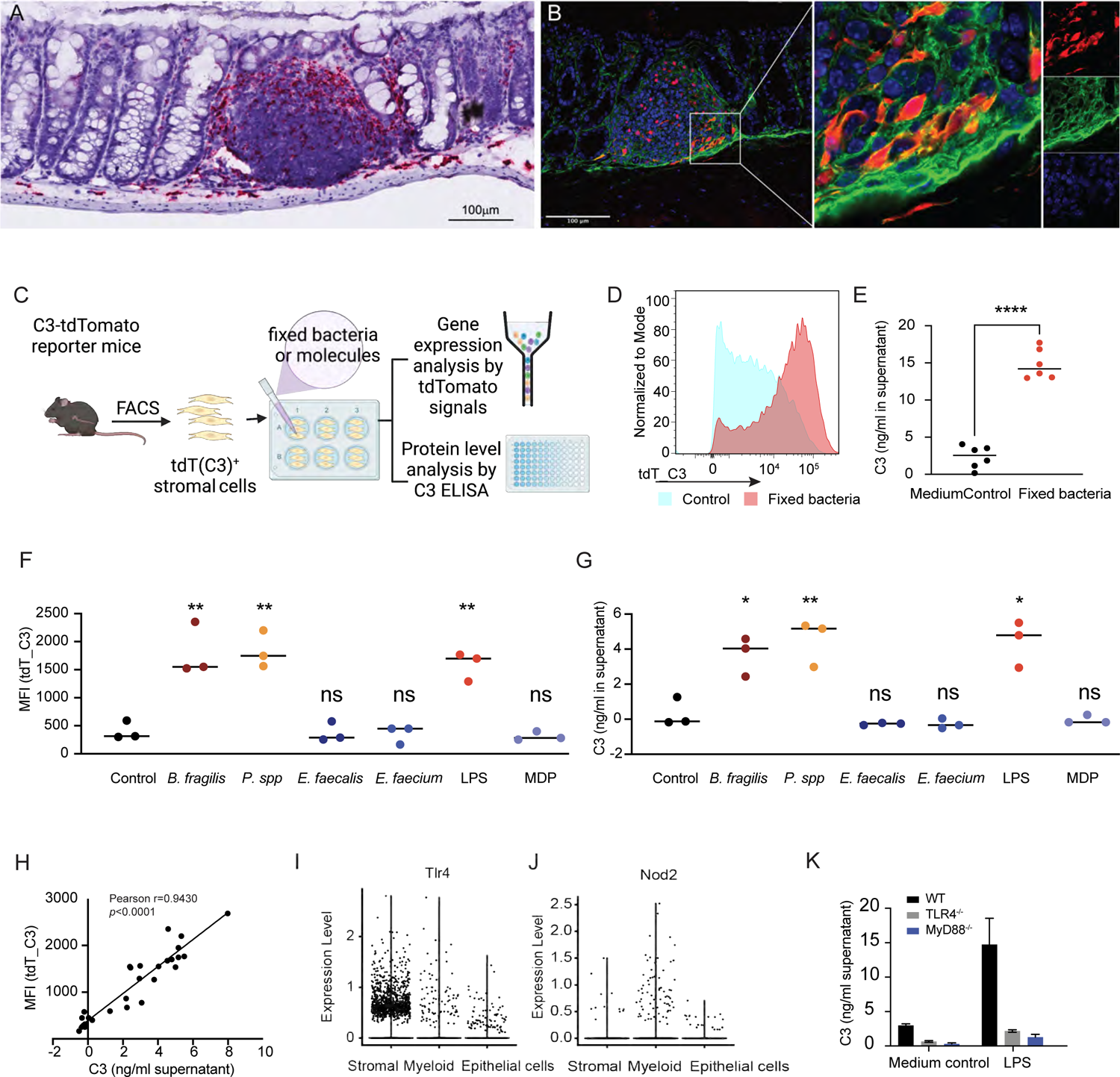
Location and function of C3-expressing intestinal stromal cells. (A). RNAscope *in situ* hybridization of C3-specific anti-sense mRNA probe (red) in mouse colon samples from WT C57BL/6 mice. (B). Immunofluorescence microscopy of colonic sections from C3^IRES-tdTomato^ reporter mice stained against tdTomato (red), PDPN (green), or DNA (blue). (C). Overall experimental scheme for the isolation of colonic C3^+^ stromal cells from C3^IRES-tdTomato^ reporter mice, which are subsequently stimulated by fixed bacteria or molecules and tested for C3 transcript expression and protein production. (D-G). Transcriptional expression levels of C3 (D and F) and C3 protein concentration in the culture medium (E and G) of colonic stromal cells stimulated by fixed bacterial strains individually or in combination. (H). Correlation of C3 transcriptional expression levels and C3 protein levels in the culture media from 4D-G. (I, J). Transcriptional expression levels of *tlr4*(I) and *nod2*(J) in the three indicated cell types from scRNASeq of C3-expressing colonic cells. (K) C3 concentration in the culture medium of colonic stromal cells from WT, TLR4^-/-^, or MyD88^-/-^ mice stimulated by LPS (1ug/ml) or medium-only as control. * p<0.05; ** p<0.01,****p<0.0001, unpaired student’s t-test.

To further test if stromal cells can produce and secrete C3 protein, we isolated colonic stromal cells from C3^IRES-tdTomato^ C57BL/6 mice by FACS and measured C3 gene expression and protein levels before and after bacterial stimulation (Figure 4C). For this, we formalin-fixed bacterial cells obtained from the feces of CR mice and used these to stimulate cultured primary stromal cells. We found increased tdTomato signal (indicating C3 transcript expression) upon bacterial exposure compared to unstimulated controls (Figure 4D). We also observed higher C3 protein levels in the extracellular medium of stromal cells 24 hours after bacterial stimulation (Figure 4E). The expression levels of tdTomato significantly correlated with protein levels of C3 in the medium (Figure 4H), confirming that this reporter system was a robust measure of C3 protein production. To further dissect the ability of bacterial strains in stimulating C3 production, we picked four representative bacterial strains: two gram-negative species, *Bacteroides fragilis* 9343 and *Prevotella* isolate CR1, the latter we reported above to increase C3 production *in vivo*, and two gram-positive species, *Enterococcus faecalis* TX0104 and *Enterococcus faecium* TX1330. We cultured them individually in their respective culture media, and then formalin-fixed them separately to stimulate stromal cell C3 production. We found both of the gram-negative bacterial strains upregulated C3 expression (Figure 4F) and C3 protein production (Figure 4G), while the gram-positive bacterial strains failed to upregulate either C3 transcript expression (Figure 4F) or protein production (Figure 4G).

To identify specific bacterial molecules participating in the induction of C3 in stromal cells, we examined expression levels of several pattern recognition receptors (PRRs) in colonic C3-expressing cells using our scRNASeq data (Figure S4D). We found that intestinal C3-expressing stromal cells express *Tlr4*, but almost no-detectable levels of *Nod2* (Figure 4I, J). Therefore, we chose two microbe-associated molecular patterns (MAMPs) to test their ability to induce C3 expression in stromal cells: lipopolysaccharide (LPS), a known TLR4 ligand from gram-negative bacteria, and muramyl dipeptide (MDP), a known NOD2 ligand from gram-positive bacteria. As expected, LPS upregulated the transcription and secretion of C3 in primary colonic stromal cells while MDP did not (Figure 4F, G). To identify innate cellular signaling pathways used by colonic stromal cells to detect LPS and trigger C3 production, we isolated stromal cells from TLR4^-/-^, MyD88^-/-^, and WT BL6 mice by FACS. Compared to WT control cells, LPS stimulation induced less C3 in both TLR4- and MyD88-knockout stromal cells indicating that the TLR4-MyD88 pathway is critical for LPS sensing and triggering of C3 production in intestinal stromal cells (Figure 4K).

### Luminal complement C3 is critical for protection against *Citrobacter rodentium* infection

To investigate the function of mucosal complement C3 on host health, we assessed the role of C3 during infection against the murine pathogen *Citrobacter rodentium*. This pathogen is the mouse counterpart to model human pathogen enteropathogenic *E. coli* (EPEC) and enterohemorrhagic *E. coli* (EHEC).^29^ We orally infected weaning-age C3-deficient mice and used age-matched WT mice as controls. C3-deficient mice were more sensitive to *C. rodentium* infection, with only a 50% survival rate, while all WT mice survived the pathogen challenge (p<0.05, Figure 5A). Adult C3-deficient adult mice also showed significantly more symptoms such as weight loss than WT mice (Figure 5B). Symptom onset occurred one week after infection and peaked five days later. We found significantly higher levels of *C. rodentium* shed in the feces of C3-deficient mice compared to WT mice at day eleven post-infection (Figure 5C). Since *C. rodentium* is known for the formation of ‘attaching and effacing’ (A/E) lesions in the colonic epithelium during infection, we examined the location of *C. rodentium* using specific rRNA FISH probes seven days after infection. There were significantly higher numbers of *C. rodentium* invasion in the epithelium of C3-deficient mice (Figure 5D) compared to WT mice (Figure 5E, 5F, p<0.005).

**Figure 5:**
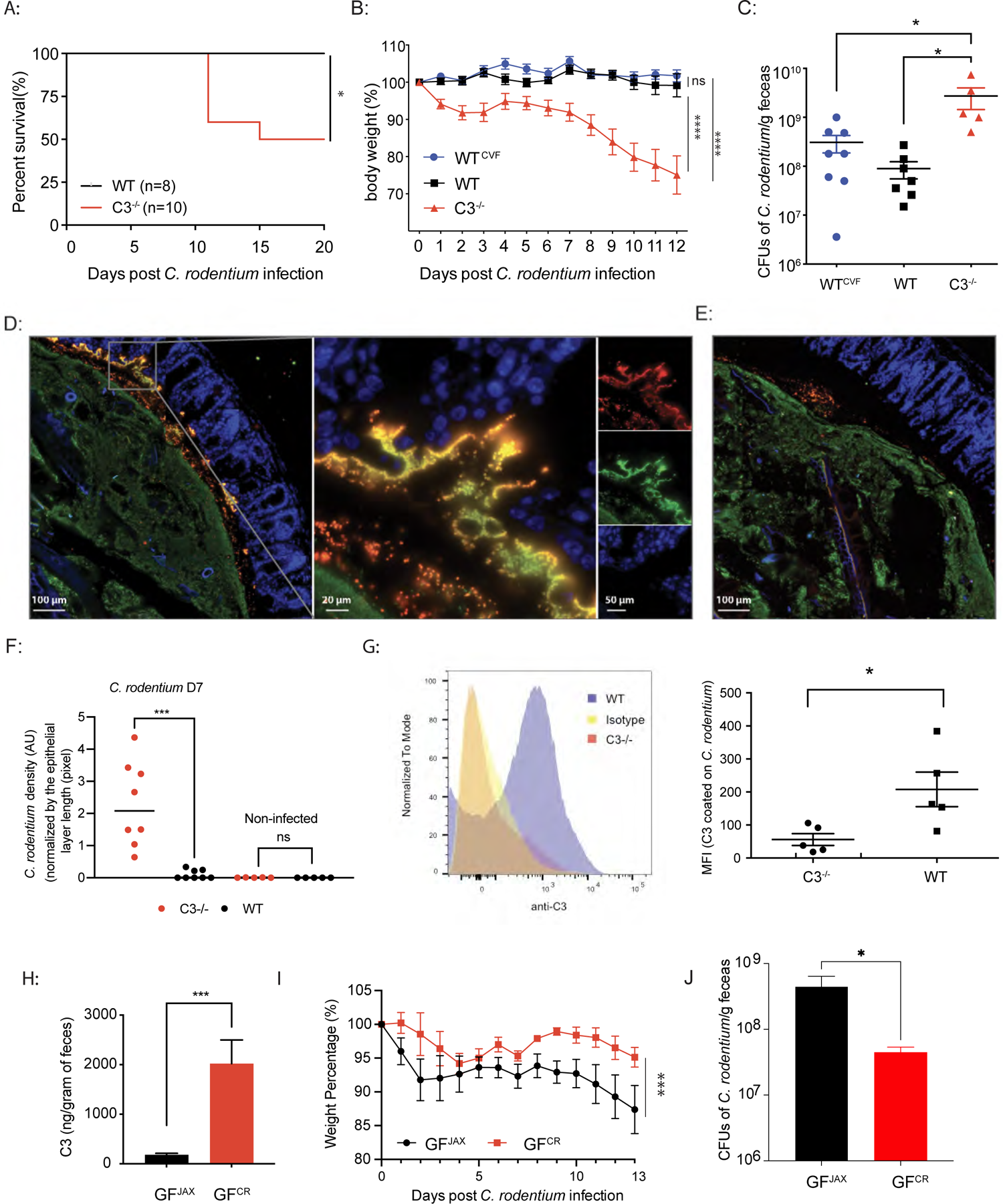
Complement factor C3 is critical for protection against *Citrobacter rodentium* infection. (A) Survival of ∼4-week-old C57BL/6 WT and C3-deficient mice following challenge with *C. rodentium (*5×10^8^ CFUs/mouse). (B). Weight loss following challenge with *C. rodentium (*5×10^8^ CFU/mouse) in three groups of ∼8-week-old mice: WT; C3-deficient; and WT depleted of serum complement with cobra venom factor (CVF). (C). Fecal CFU counts of *C. rodentium* at day 11 post-infection in untreated WT, WT treated with CVF (WT^CVF^), or C3-deficient mice. (D, E). Images of cross-section of colons containing fecal matter from *C. rodentium*-infected mice that are C3-deficient (D) or WT €. Samples were stained with *C. rodentium* specific rRNA FISH probes (red), Eub338 rRNA FISH probes (green), and DAPI (blue). (F). *C. rodentium* density under the epithelial layer of C3-deficient or WT mice, normalized by colon length (n = 8). (G). Flow cytometry histogram and quantification of C3 protein on GFP^+^ *C. rodentium* at day 7 post-infection. (H). Fecal C3 levels of gnotobiotic mice two weeks after colonization with JAX or CR microbiota. (I). Weight loss of gnotobiotic mice colonized with JAX or CR microbiota following challenge with *C. rodentium* (10^10^ CFUs/mouse). (J). Fecal CFU enumeration of *C. rodentium* on day eleven post-infection in gnotobiotic mice colonized prior to challenge with JAX or CR microbiota. ns=not significant, *<0.05; ** p<0.01,***p<0.001,Student’s t-test

To determine whether mucosal C3 was deposited on bacterial cells during infection, we infected mice with *C. rodentium* LB1, a strain which constitutively expresses GFP,^30^ and performed flow-cytometric analysis on bacterial cells in fecal samples from infected WT and C3-deficient mice. Immunostaining of GFP^+^ *C. rodentium* cells with fluorescent anti-C3 antibody showed that the microbes were coated with C3 at day seven post-infection (Figure 5G). To rule out whether the classical or lectin pathways are required for this phenotype, we infected C1qa- and MBL-deficient mice with GFP+ *C. rodentium*, we found similar levels of fecal C3 and C3-coated GFP^+^ *C. rodentium* cells (Figure S5A-C), suggesting that the deposition of C3 does not rely on the classical or lectin pathway. This was consistent with the report that C1q is not essential for protection against *C. rodentium* infection.^27^

To confirm the role of luminal C3, as opposed to serum C3, in *C. rodentium* infection, we treated SPF mice with cobra venom factor (CVF) to specifically deplete plasma-circulating C3 without depleting gut luminal C3 (Figure S5D, E). Subsequently, mice were challenged with *C. rodentium* and monitored for symptoms. We found CVF treatment did not compromise the resistance to *C. rodentium* infection displayed by WT mice. There was no significant difference in weight loss (Figure 5B) or bacterial burden in CVF treated compared to non-CVF treated WT controls (Figure 5C). These results indicate that mucosal complement C3 is sufficient to confer resistance to this enteric pathogen independently of C3 production at non-mucosal sites.

We next tested if modulation of luminal C3 by different gut microbiota could affect host susceptibility to *C. rodentium* challenge. We colonized GF mice with JAX or CR microbiota resulting in GF^JAX^ with low luminal C3 level or GF^CR^ with high luminal C3 level, respectively (Figure 5H). When mice were challenged with a high dose of *C. rodentium* (2X10^10^ CFU/mouse), we found that GF^JAX^ mice lost significantly more weight during the infection compared GF^CR^ mice (Figure 5I). GF^JAX^ mice also shed significantly higher numbers of *C. rodentium* in their feces compared to GF^CR^ mice on the 11^th^ day after infection (Figure 5J). These results demonstrated that the level of luminal C3 driven by the unique compositions of the host gut microbiota modulated the degree of susceptibility to enteric infection by *C. rodentium*.

Our results with the murine enteric pathogen *C. rodentium* suggest that mucosal C3 can opsonize select enteric pathogens during enteric infections and that this can occur independently of the classical or lectin pathways. Variations in luminal C3 levels due to the host commensal microbiota can impact the host’s susceptibility to enteric pathogen infection.

### Luminal C3 production increases significantly during enteric infection

We reasoned that if luminal C3 levels play a major role in controlling enteric infections, *C. rodentium* infection may lead to up-regulation of mucosal C3 at the transcriptional and protein levels. To test this hypothesis, C3^IRES-tdTomato^ reporter mice were infected with *C. rodentium*; transcriptional levels of C3 were measured by flow cytometry and fecal C3 protein levels were measured by ELISA (Figure 6A). We found that fecal C3 protein levels increased significantly during infection (Figure 6B), which correlated with C3 gene transcriptional expression as measured by the total tdTomato signal in cells from the colonic subepithelial compartment (Figure 6C). Gating on tdTomato-positive cells and following select genetic markers for various cell types, we found that stromal cells, CD45^+^ cells, and IECs are also major sources of C3 production during infection (Figure S6A, B).

**Figure 6:**
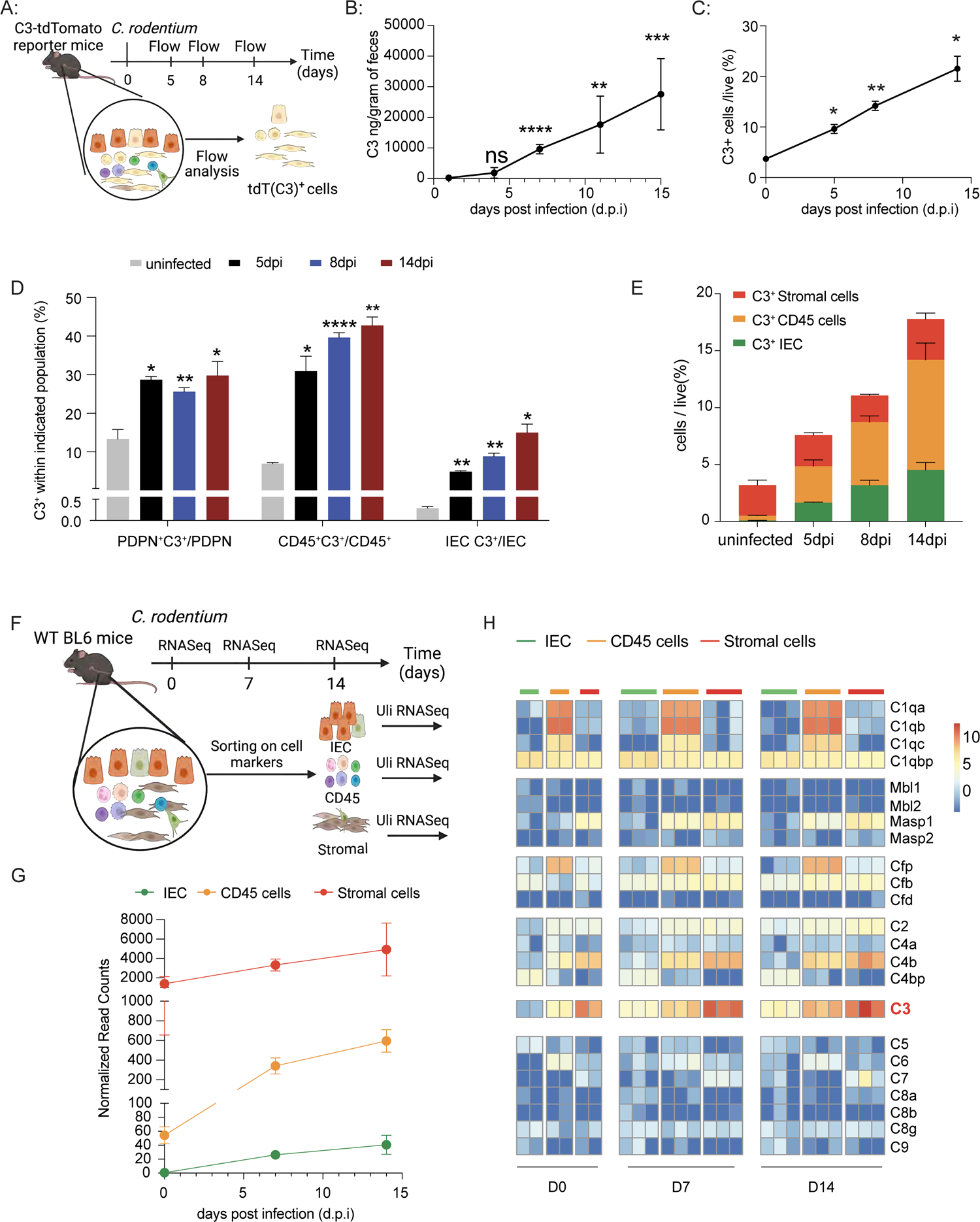
Fecal levels of Complement C3 increase significantly during infection. (A). Overall experimental timeline of flow cytometric analysis of C3-expressing cells in C3^IRES-tdTomato^ reporter mice after *C. rodentium* infection. (B). Fecal C3 protein levels at 1, 4, 7, 11 and 15 days post-infection (dpi). (C). Percentage of live cells isolated from the colonic lamina propria of C3^IRES-tdTomato^ reporter mice transcriptionally expressing C3 on days 0, 5, 8 and 14 post-infection. (D). Percentage of cells transcribing C3 among IECs, stromal, and CD45^+^ cells. (E). Percentage of IEC, stromal, or CD45^+^ cells transcribing C3 amongst the total live cell population isolated from colonic lamina propria at indicated days post-infection. (F). Overall experimental timeline of RNASeq analysis of sorted cell populations in WT C57BL/6 mice after *C. rodentium* infection. (G). RNA-seq analysis of C3 gene expression levels in the three indicated cell populations in WT C57BL/6 mice. (H). Heatmap of complement system gene expression levels in the three indicated cell populations of WT C57BL/6 mice. ns: not-significant, ** p<0.01, *** p<0.001, **** p<0.0001 Student’s t-test

Compared to uninfected controls, mice infected with *C. rodentium* had an increased proportion of stromal cells transcriptionally expressing C3 in the total colonic stromal cell population (30% versus 12%), which coincided with the early phase of infection as *C. rodentium* begins to attach to enterocytes.^31^ The percentage of stromal cells transcribing C3 was maintained at ∼30% throughout the infection. CD45^+^ cells transcribing C3 increased from 5% to ∼30% of total CD45^+^ cells by day five post-infection and reached a level of ∼40% by day eight. This level of expression was maintained in CD45^+^ cells through day 14. IECs transcribing C3 increased from ∼0.5% to 5% of total IECs by day five and reached a level of ∼10% by day eight which continued to rise to ∼15% through day 14 (Figure 6D). In terms of cell numbers, due to the large number of total IECs in the colon, IEC hyperplasia, and the infiltration of CD45^+^ cells into the gut during infection, the composition of the C3-transcribing population changed from predominantly stromal cells to predominantly CD45^+^ and IECs as the infection progressed (Figure 6E, Figure S6B).

We next performed bulk RNA-Seq on sorted IECs, CD45^+^, and stromal cells from uninfected and infected C57BL/6 mice to confirm our ELISA and reporter assay results (Figure 6F). We found significant increases in C3 gene expression in all three populations at day seven and fourteen post-infection (Figure 6G), suggesting all contributed to C3 transcriptional expression. Nonetheless, stromal cells still showed the highest C3 mRNA expression levels during infection when compared to other host cell populations (Figure 6G), consistent with the results in reporter mice.

We also looked for increased transcript expression of other complement system proteins using bulk RNA-Seq and compared infected and non-infected controls of the same three cell populations in WT C57BL/6 mice. We found that in addition to C3, both Cfb and C4b are significantly upregulated in stromal and CD45^+^ cells. However, the majority of genes encoding MAC components (C5-C9) were not expressed at detectable levels by RNA-Seq in any of the three sorted populations (Figure 6H), which is consistent with low C5 protein levels in WT fecal specimens from day eleven post-infection (Figure S6C). Pendse *et al.* reported that gut transit time was unaltered in C3-deficient mice^27^, which suggested that the differences between WT and C3-deficient mice were not due to changes in intestinal motility.

### C3-dependent neutrophil-mediated phagocytosis is critical for protection against *C. rodentium* infection

We reported above (Figure 5C) that C3-deficient mice had significantly greater organism loads during infection than C3-sufficient mice. Thus, we wanted to determine the major mechanisms used by the gut immune system to limit organism load. Since components of the MAC (C5-C9) are not upregulated or secreted by C3-producing cells in the colon during infection, we examined the role of opsonophagocytosis in controlling *C. rodentium* infection. Flow cytometric analysis of intestinal cells isolated from C3^IRES-tdTomato^ reporter mice showed a population within CD45^+^ cells that incrementally upregulated C3 expression throughout the course of infection (Figure 7A). While in a homeostatic environment only ∼4% of C3-expressing CD45^+^ cells express high C3 levels, this population rose to ∼50% 14 days post-infection (Figure 7B). Further analysis of this high-C3-expressing population revealed they were myeloid cells (CD11b^+^), with the majority being neutrophils, followed by macrophages and monocytes (Figure 7C, Figure S7A, B).

**Figure 7:**
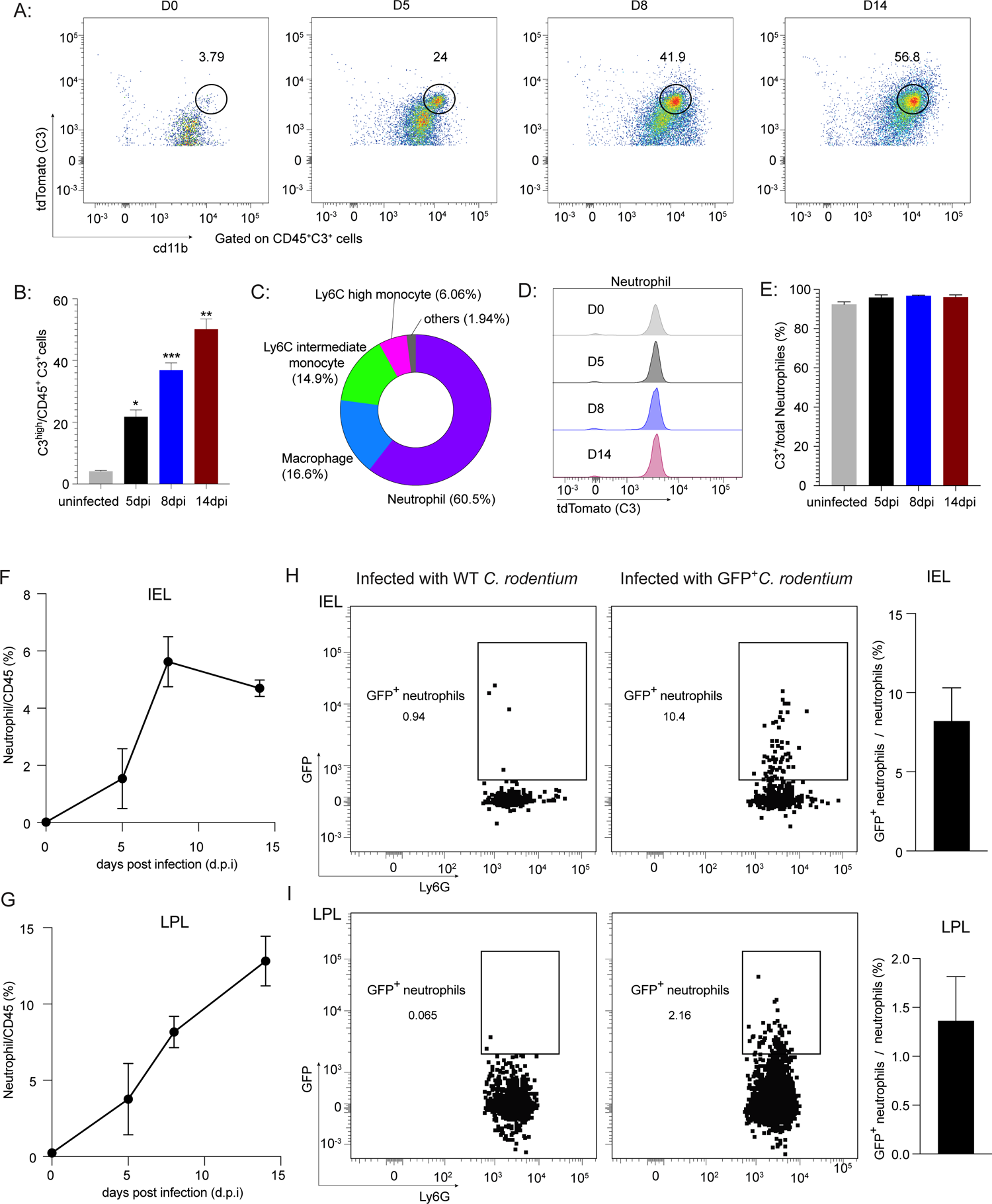
C3-mediated neutrophil phagocytosis is crucial for clearing *Citrobacter rodentium* during enteric infection. (A-B). Representative gating (A) and frequencies (B) of the high-C3-expressing subset in colonic subepithelial CD45^+^C3^+^ cells during homeostasis (D0) and at 5, 8 and 14 days post *C. rodentium* infection. (C). Percentage of neutrophils, macrophages, Ly6C-intermediate and Ly6C-high monocytes in the high-C3-expressing CD45^+^ cells at 14 days post *C. rodentium* infection. (D). Representative histograms of C3 gene expression (MFI in C3-tdTomato reporter mice) in neutrophils at 0, 5, 8, and 14 days post-infection (dpi) in C3^IRES-tdTomato^ reporter mice. (E). Percentage of the C3-expressing neutrophils amongst total neutrophils from colonic lamina propria at indicated days post-infection. (F). Percentage of neutrophils in CD45^+^ cells in epithelium (F) or colonic lamina propria (G) at day 0, 5, 8, and 14 post-infection. (H-I). Representative gating and frequency of GFP^+^ *C. rodentium* uptake in neutrophils in the colonic intraepithelial (H) and lamina propria (I) layers at seven days post-infection. Left panels are the mice infected with WT *C. rodentium* as control for GFP^+^ gating. Right panels are the mice infected with GFP^+^ isogenic *C. rodentium*. * p<0.05,** p<0.01, *** p<0.001, Student’s t-test

We then analyzed C3 expression dynamics in these three myeloid populations during infection. While only a subset of macrophages and monocytes expressed C3 (Figure S7C-E), neutrophils uniformly expressed high levels of C3 (Figure 7D, E). Among the myeloid cell population, neutrophils had the highest tdTomato Mean Fluorescence Intensity (MFI) and maintained this level throughout the infection (Figure 7D, Figure S7C-E). We found that neutrophils were robustly recruited to the colonic lamina propria and epithelial layer during infection (Figure 7F, G) where they became the major source of high-C3-expressing CD45^+^ cells (Figure 7C, Figure S7B). The dominant presence and activity of neutrophils is consistent with the concept that phagocytosis of microbes is imperative for C3-dependent bacterial clearance.

To explore the hypothesis that mucosal C3 was acting as an opsonin to enhance bacterial phagocytosis, we measured the magnitude of bacterial uptake by neutrophils in the mucosal environment. WT mice were infected with GFP^+^ *C. rodentium*, and neutrophils from the mucosal epithelial and lamina propria layers were collected and analyzed by flow cytometry seven days after infection. We found a significant fraction of GFP^+^ neutrophils in both layers (Figure 7H, I), suggesting that neutrophils had phagocytosed GFP^+^ *C. rodentium* within the host mucosal environment.

Finally, we wanted to determine whether gut complement was important to the clearance of other enteric pathogens. Therefore, we tested the human pathogen EHEC in a murine environment. We infected 6-week-old C3-deficient or WT C57BL/6 mice with GFP^+^ EHEC strain EDL931 via the oral gastric route. We found that adult C3-deficient mice were highly susceptible to EHEC infection, showing only a 16% survival rate. In contrast, WT mice were substantially more resistant to EHEC infection with only one out of eleven mice succumbing to the infection (Figure S7G). We collected fecal samples from WT and C3-deficicent mice at four days post-infection and performed flow cytometric analysis using an anti-C3 antibody to assess C3 opsonization of EHEC. C3 was detected on fecal GFP^+^ EHEC EDL931, demonstrating that EHEC can also be significantly opsonized by C3 in the lumen of WT mice (Figure S7H).

## Discussion

The complement cascade is an evolutionarily conserved part of our immune system. For decades, complement has been considered primarily a hepatocyte-derived and serum-effective system responsible for defending against the invasion of microbes into host tissues. However, initial host-pathogen interactions often occur at the apical side of epithelial barriers, and the role of complement in these surfaces has not been thoroughly explored. In this study, we found that local colonic cell populations can sense gut commensal microbes and produce a substantial quantity of complement C3 in the gut lumen during homeostasis. The level of luminal C3 is regulated by the composition of the microbiota, which confers individualized luminal C3 levels. Furthermore, luminal C3 increases significantly during enteric infection with the murine pathogen *C. rodentium*, where C3 is required for protection against this and EHEC strain EDL931, an important human diarrheagenic enteropathogen. Even though pathogenic *E. coli* infections are a major worldwide cause of diarrheal disease, especially in children, the mechanisms for host defense are not well understood. *C. rodentium* is considered a murine A/E pathogen and serves as a mouse model for EPEC and EHEC infections, while EHEC also produce Shiga-like toxins.^32^ Our study expands our understanding of the role of the microbiota in shaping variability in host infection susceptibility and severity and reinforces the important concept that the microbiota can indirectly influence a host’s response to infection by priming immune defense mechanisms before a pathogen is encountered.

Diet is a major factor that shapes the composition of the gut microbiota. In low- and middle-income countries, malnutrition and undernutrition are major public health concerns.^33^ Malnourished children exhibit delayed maturation of the microbiome and have greater susceptibility to enteric pathogens.^34, 35^ Malnutrition and enteric disease susceptibility may be both a cause and a consequence of each other. Pathogenic *E. coli* (including EHEC and EPEC) are thought to be the most common enteric pathogens found to be associated with diarrheal disease in low- and middle-income countries. Our findings may directly relate disease susceptibility and severity to alterations in the microbiota resulting from undernutrition.

Fecal C3 was previously connected to non-infectious diseases. Local gut complement C3 deposition was observed on the surface of the colonic epithelium in biopsies or resection specimens from IBD patients.^36^ However, due to inflammation in IBD patients and increased vascular permeability causing unpredictable extravasation of serum proteins, it could not be determined whether C3 was synthesized locally in the gut or secreted into the lumen during homeostasis. Here, we provide direct evidence that C3 can be locally transcribed by three different cell populations in the steady-state using C3-reporter mice, scRNASeq, flow cytometry, and validation by ultra-low input RNA-Seq on sorted cell populations. We found that C3 protein exists in the lumen of healthy mice and humans and is regulated by the gut microbiota. There was a local increase in C3 transcription levels in stromal, epithelial, and myeloid cells and an increase in C3 protein levels in the lumen during enteropathogenic infection. Together, these data revealed the dynamic local complement C3 response in the gut.

The finding that stromal cells, (fibroblasts in particular) are the major source of C3 in both mice and humans during homeostasis is an unexpected finding. Traditionally, fibroblasts are non-hematopoietic (CD45^-^), non-epithelial (Epcam^-^), and non-endothelial (CD31^-^) structural cells that provide essential connective tissue (stroma).^37^ They define the architecture of the tissue microenvironment by depositing and remodeling extracellular matrix (ECM) components. Recent studies have uncovered important functions for these cells in initiating, supporting, or suppressing innate and adaptive immune responses in lymphoid and non-lymphoid tissues.^38^ One such example is the role of fibroblasts in the tumor microenvironment (TME). Tumor-associated fibroblasts are abundant in the TME and radically influence disease progression and metastasis across many different cancers.^39–43^ Here, we established a novel function of stromal cells to defend against pathogens through their expression of complement C3. C3-expressing fibroblasts are primarily located in colonic LFs, which are comprised of B-, T-, myeloid, and supporting cells including fibroblasts. We have demonstrated that these fibroblasts can directly sense bacterial signals *in vitro* and respond to these cues by producing C3.

Our study on intestinal C3 opens a new avenue of research for local complement systems beyond synthesis in the liver and activity in blood. We explored transcriptional profiles of central components of the complement system using scRNASeq and ultra-low input RNA-Seq on sorted cell populations, as well as C3 and C5 protein levels during homeostasis and infection. Two dominant cell types were involved in gut complement production: stromal cells were the dominant source of intestinal C3, while myeloid cells were the dominant source of gut C1q. These findings were validated by scRNASeq data from healthy human gut tissues.^26^ The difference in cellular sources for C1q and C3 is notable as the complement cascade is currently viewed as an integrated, interdependent system. In our study, we found gut luminal C3 recognizes and binds to pathogens and is required for phagocyte-mediated killing of these microbes; this is consistent with the classical understanding of complement. Other research groups have explored the effects of local complement systems in assessments relevant to our study. C1q is not required for immune defense in the intestine but rather plays a critical role in regulating gut motility by interacting with the enteric nervous system.^27^ Additional studies also show that the complement system plays an important role in the ocular mucosa. During infection with ocular pathogens such as herpes simplex virus type 1 (HSV-1), C3d is deposited intracellularly in the corneal epithelium of vaccinated animals to confer protection, which is absent in C3-deficient mice and associated with a delay in viral clearance in vaccinated C3-deficient animals.^44^ Together, our study and others’ show complement system components may play different roles locally than they do systemically. While the functions of these components will need to be further clarified, our study raises questions about the role of complement components in other mucosal surfaces of the body including the respiratory and urinary tracts.

## Acknowledgments

We thank the staff in our animal facility for their support in animal husbandry, and all Kasper laboratory members for their comments and support. We thank Ian Magill, Niket Patel, Gang Wang, Yutao Wang, Teshika Jayewickreme, and Juliana Lee for their help with flow cytometry analysis, scRNASeq, and UltraLowInput (Uli) RNASeq. We thank Dr. Philip Ahern (Cleveland Clinic) for advice and discussions. We thank Dr. Claudia Kemper (National Institute of Health) for C3^IRES-tdTomato^ reporter mice. We thank Dr. Lynn Bry (Brigham and Women’s Hospital) for *C. rodentium* DBS100 and GFP^+^ *C. rodetium* LB1. We thank Harvard Gnotobiotic Core, Harvard Immunology Flow Cytometry Core, Microscopy Resources on the North Quad (MicRoN) Core, Harvard Rodent Histopathology Core, and Histology Core at BIDMC for their help and support. This work was supported in part by NIH, NIAID 5R01AI148273 to D.L.K., and in part by NIH R01AI157106 and R35GM124724 to A.H.

## Author contributions

Planning and conceptualization, M.W, W.Z., and D.L.K; flow cytometry, sample collection, and ELISAs, M.W., W.Z., X.S., B.B, T.Y., H.S.C., F.G., M.I., D.C. and C.T.; mouse microbiome work and analysis, M.W.; human microbiome work and analysis: M.W., R. L, J.M., E.D., and A.H. imaging, M.W., W.Z., Y.W, D.R., D.Y., W.A. and S.H.; scRNASeq, UliRNASeq and computational analysis, M.W.; provision of key resources, S.H., I.M.C, J.R.M., A.H., J.J.M., C.B. and D.L.K; Writing and Editing Manuscript, M.W, W.Z., A.H., J.J.M and D.L.K with help from all authors.

## Declaration of interests

J.R.M is a co-founder, stakeholder, and advisor for Vizgen, Inc.

## Lead Contact and Materials Availability

Please direct requests for resources and reagents to Lead Contact, Dennis Kasper. Raw 16S rRNA gene sequencing are deposited to NCBI under BioProject XXX, scRNASeq and UliRNASeq data are deposited to NCBI under GSEXXX.

## Supplementary Figures and Tables

**Figure S1:**
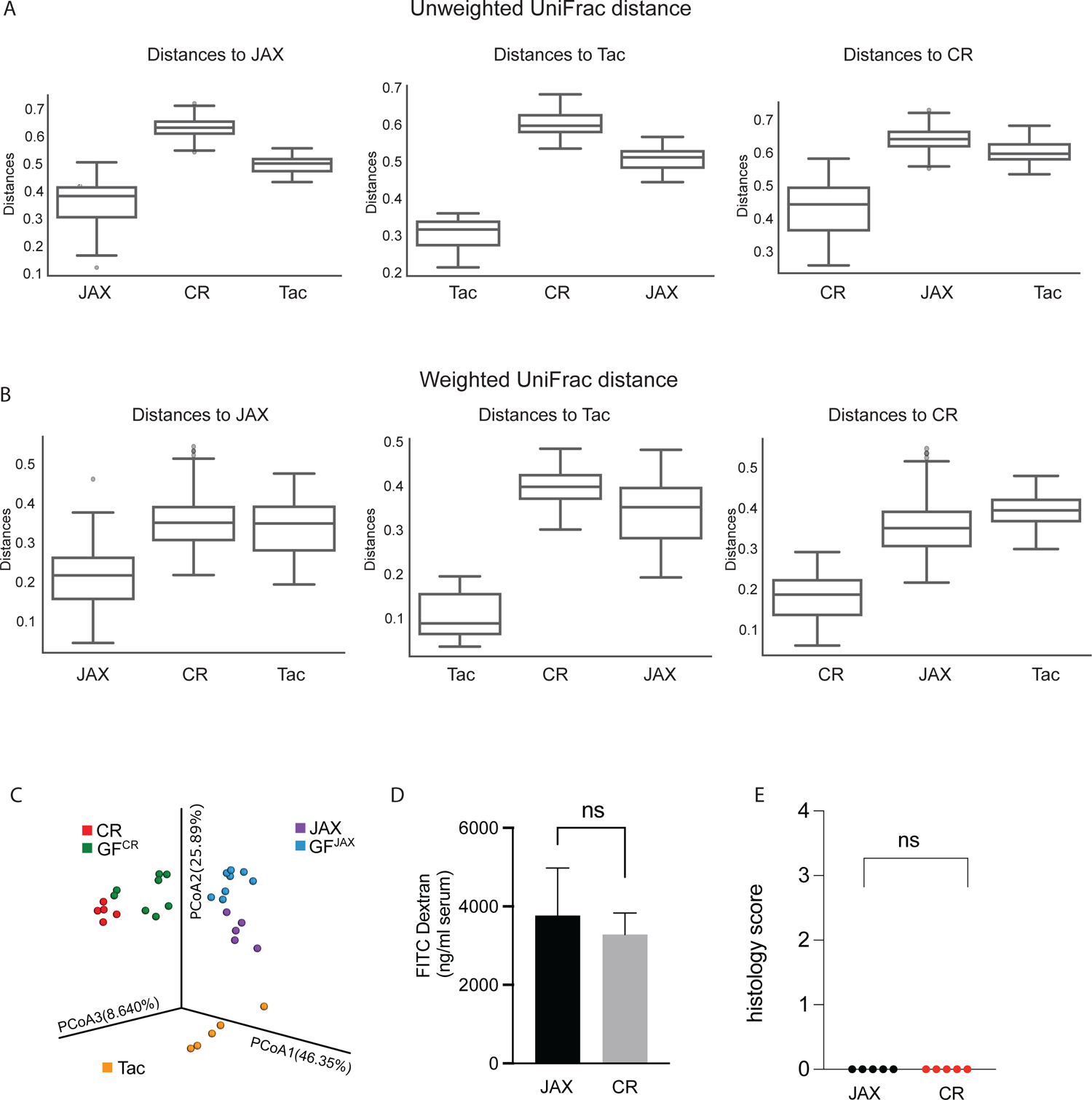
Microbiome analysis of fecal samples and gut barrier function assays in C57BL/6 mice from three commercial vendors. (A, B). Pairwise unweighted UniFrac distances (A) and weighted UniFrac distances (B) of fecal bacterial community composition within and across three commercial vendors: Jackson (JAX); Taconic (Tac); and Charles River (CR). (C) Principal Coordinates Analysis (PCoA) of weighted UniFrac distance measurements based on the 16S rRNA gene sequencing data of the fecal bacterial composition in samples collected from C57BL/6 JAX, TAC, CR, or gnotobiotic mice colonized with JAX (GF^JAX^) or CR microbiota (GF^CR^) ((q-values=0.001, PERMANOVA). (D). Fluorescence measurement of serum collected from JAX and CR mice orally gavaged with FITC-dextran. (E). Histology scoring of colons collected from JAX and CR mice. ns: not-significant, unpaired student’s t-test.

**Figure S2:**
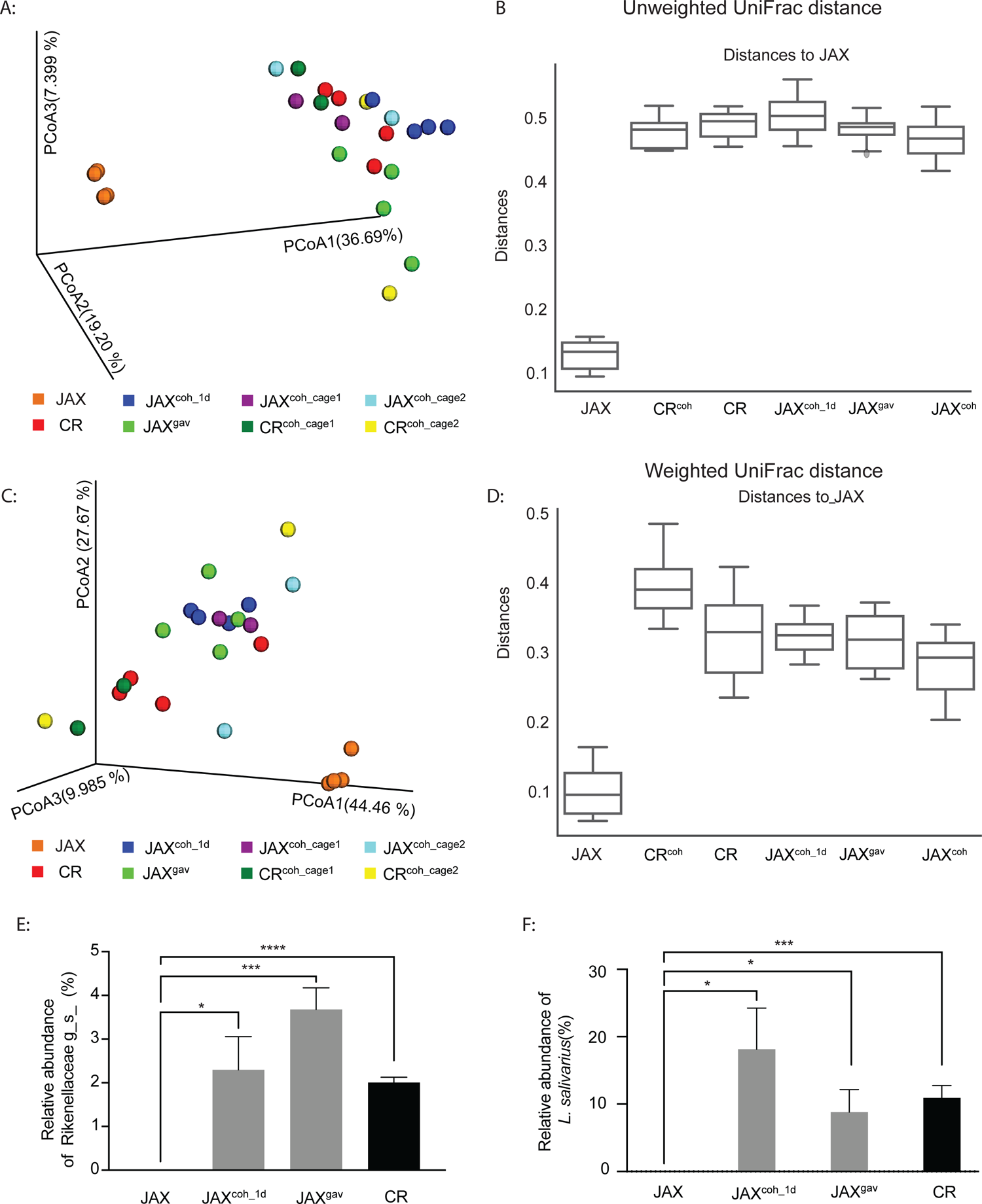
Microbiome analysis of fecal samples in co-housing experiments. (A,C) Principal coordinates analysis (PCoA) of unweighted UniFrac distance (A) and weighted UniFrac distance (C) measurements based on the 16S rRNA gene sequencing data of fecal bacterial composition in samples collected from mice as indicated (q-values=0.001 between JAX to all other groups, PERMANOVA). (B, D) Pairwise unweighted UniFrac distances (B) and weighted UniFrac distances (D) of the composition of bacterial communities in fecal samples within JAX and across groups compared to JAX. (E, F) Relative abundance of *Rikenellaceae spp.* and *L. salivarius* in JAX or CR mice, as well as in JAX mice cohoused with CR mice for 1 day (JAX^coh_1d^), or in JAX mice gavaged with CR microbiota (JAX^gav^). ns: not-significant,*p<0.05, *** p<0.001, **** p<0.0001, unpaired student’s t-test.

**Figure S3:**
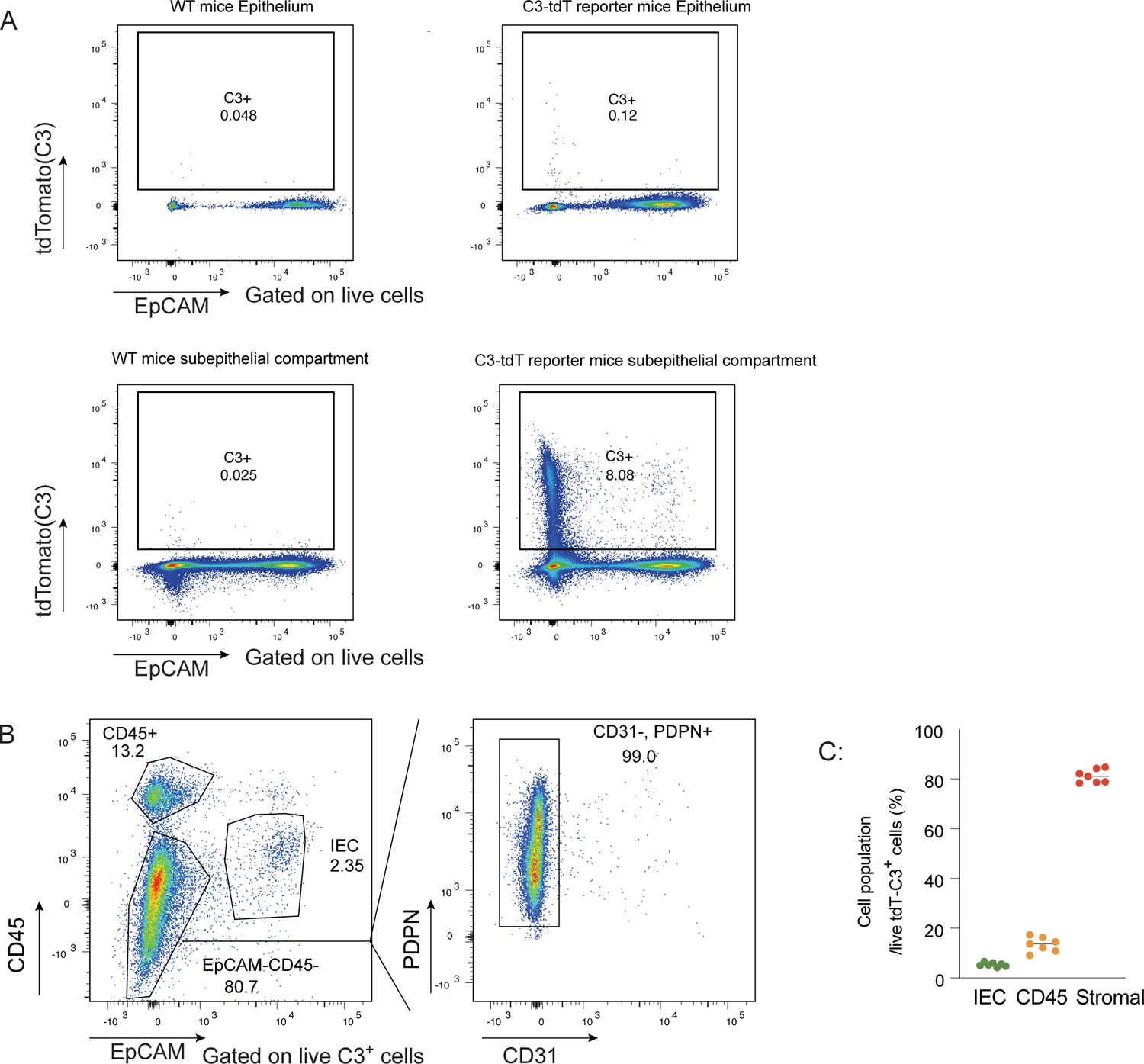
Flow cytometric analysis of C3-expressing colonic cells in C3^IRES-tdTomato^ reporter mice. (A) Flow cytometry gating of live colonic epithelial and subepithelial compartment of C3^IRES-tdTomato^ reporter mice showing the frequency of C3-tdTomato^+^ cells with WT mice as control. (B) Flow cytometry gating of live C3^+^ colonic cells in subepithelial compartment of C3^IRES-tdTomato^ reporter mice showing the frequencies of CD45^+^, IEC, CD45^-^EpCAM^-^, and CD31^-^PDPN^+^ cells. (C) Percentage of IECs, stromal, or CD45^+^ cells in total colonic C3-expressing cells in the subepithelial compartment as assessed by flow cytometry in (B).

**Figure S4.**
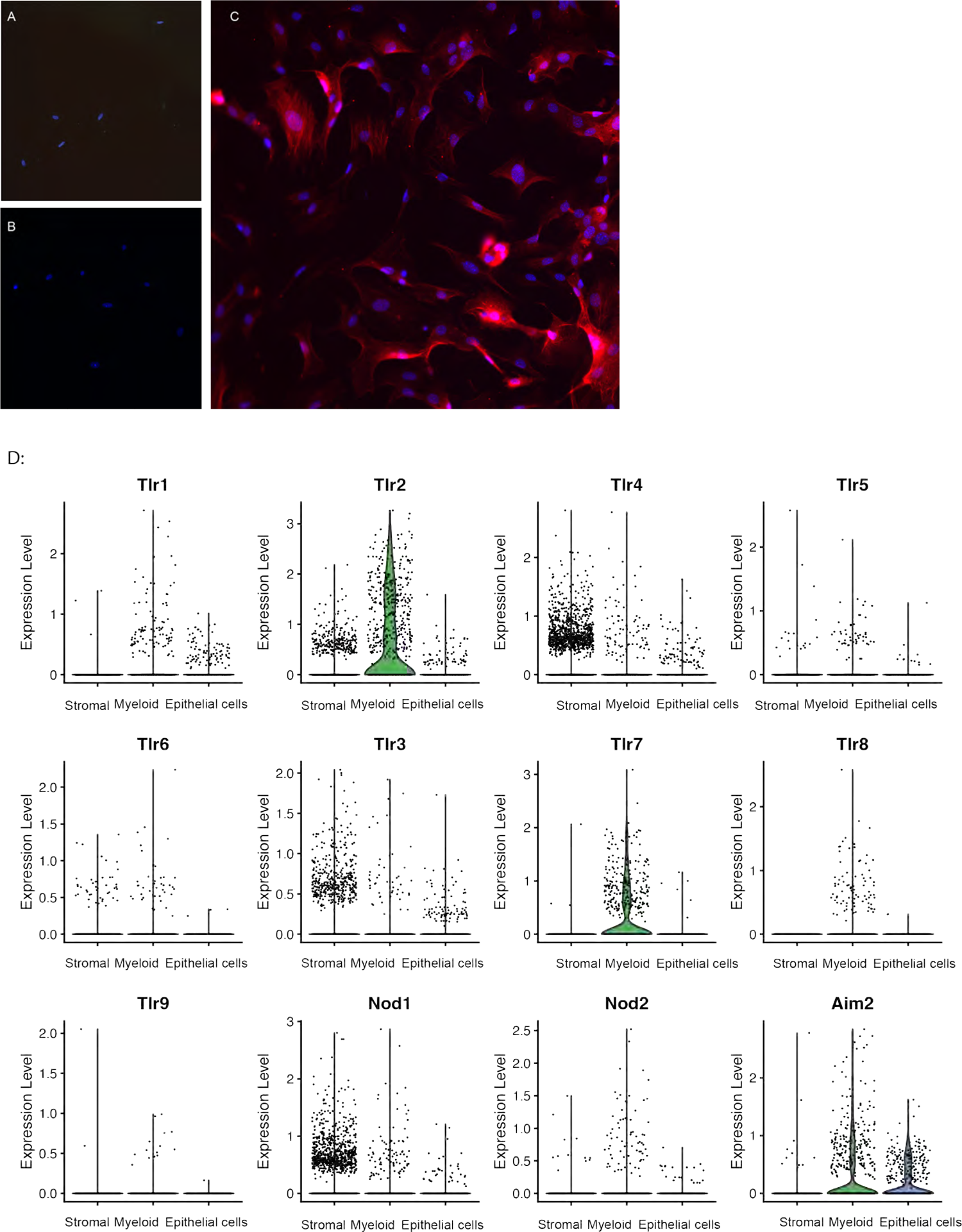
Immunofluorescence microscopy of C3-expressing fibroblasts and transcriptional expression levels of receptor genes in C3-expressing colonic cells. (A-C) Immunofluorescence microscopy of C3-expressing fibroblasts isolated from the colon of C3^IRES-tdTomato^ reporter mice, stained with a primary antibody against tdTomato (A), a secondary antibody against tdTomato (B), or both primary and secondary antibodies (red) against tdTomato (C), as well as Hoechst (blue). (D) Violin plot showing indicated receptor gene expression levels in C3-expressing stromal, myeloid, and epithelial cells identified in Figure 3C from scRNASeq of tdTomato^+^ mouse colonic cells using C3^IRES-tdTomato^ reporter mice.

**Figure S5.**
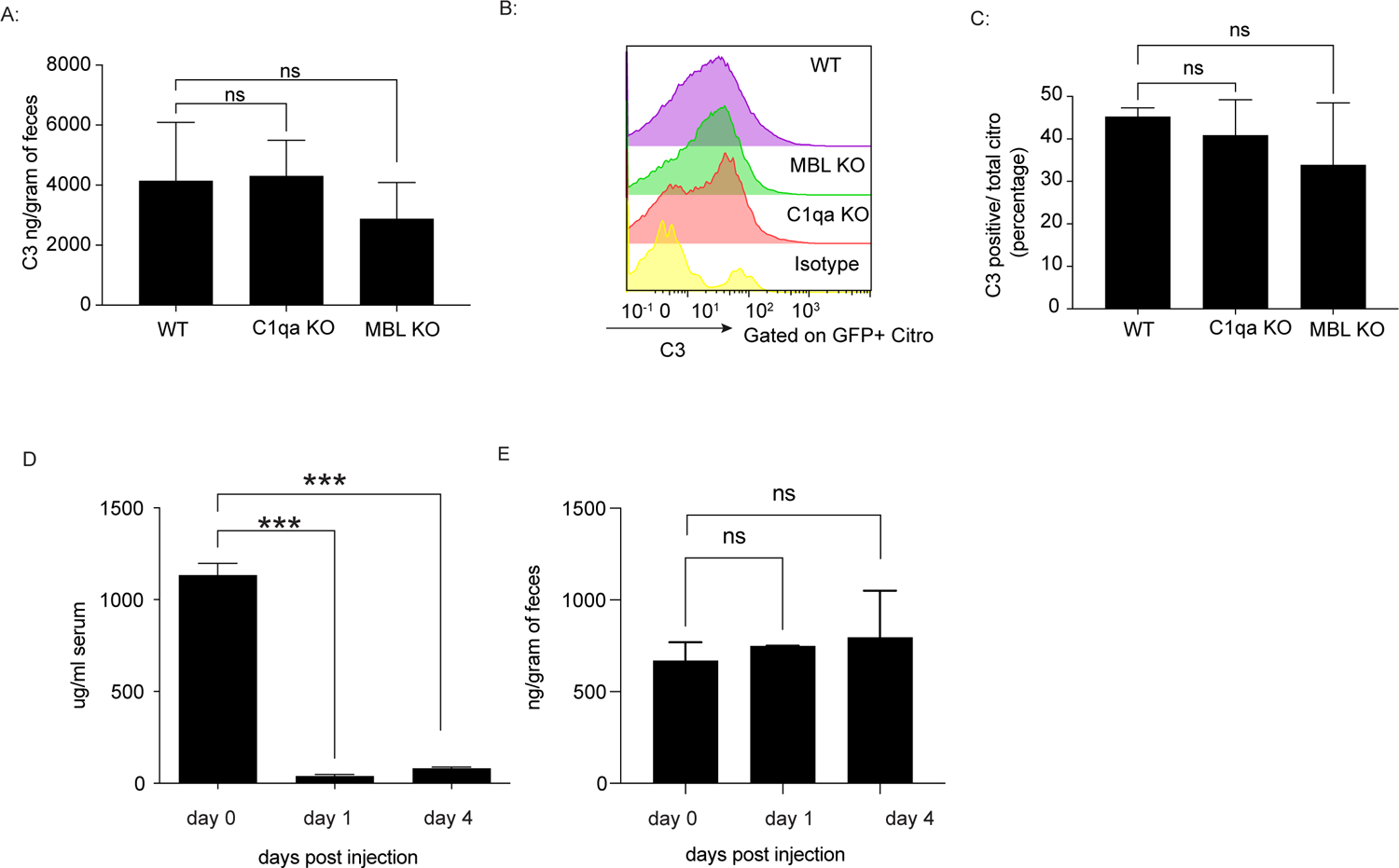
C3 levels and activity in C1qa deficient, MBL deficient mice, and WT mice treated with cobra venom factor. (**A**) Fecal C3 levels at day seven post-*C. rodentium*-infection in C1qa-deficient or MBL-deficient mice with WT mice as control. (**B-C**) C3 opsonization of GFP^+^ *C. rodentium* at day seven post infection in WT, MBL-deficient, or C1qa-deficient mice by flow cytometric analysis using anti-C3 antibody or isotype control (**B**) and the frequency of C3^+^ *C. rodentium* (**C**). (**D, E**) Serum C3 levels (D) and fecal C3 levels (E) in WT mice treated with cobra venom factor at day 0, 1, and 4 after treatment.

**Figure S6:**
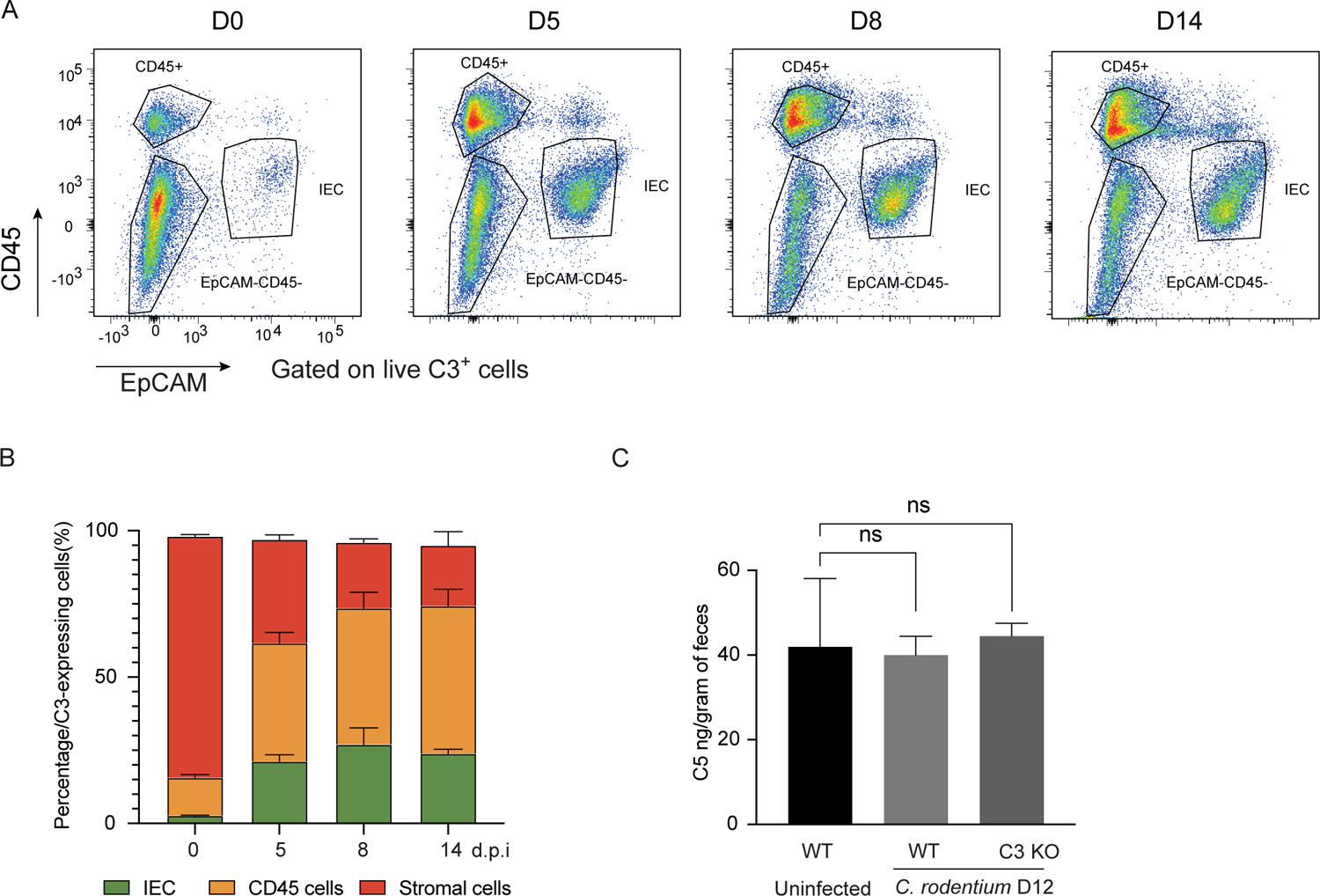
Composition of total colonic C3-expressing cells and fecal C5 levels during *C. rodentium* infection. **(A)** Flow cytometry gating of live C3^+^ colonic cells in subepithelial compartment of C3^IRES-tdTomato^ reporter mice showing the frequency of CD45^+^, IEC, or CD45^-^EpCAM^-^ cells. **(B)** Percentages of IEC, CD45^+^, or stromal cells in live C3^+^ colonic cells in the subepithelial compartment of C3^IRES-tdTomato^ reporter mice during *C. rodentium* infection. (C). Fecal C5 levels in uninfected WT mice or infected WT and C3-deficient mice at day 12 post-*C. rodentium* infection.

**Figure S7:**
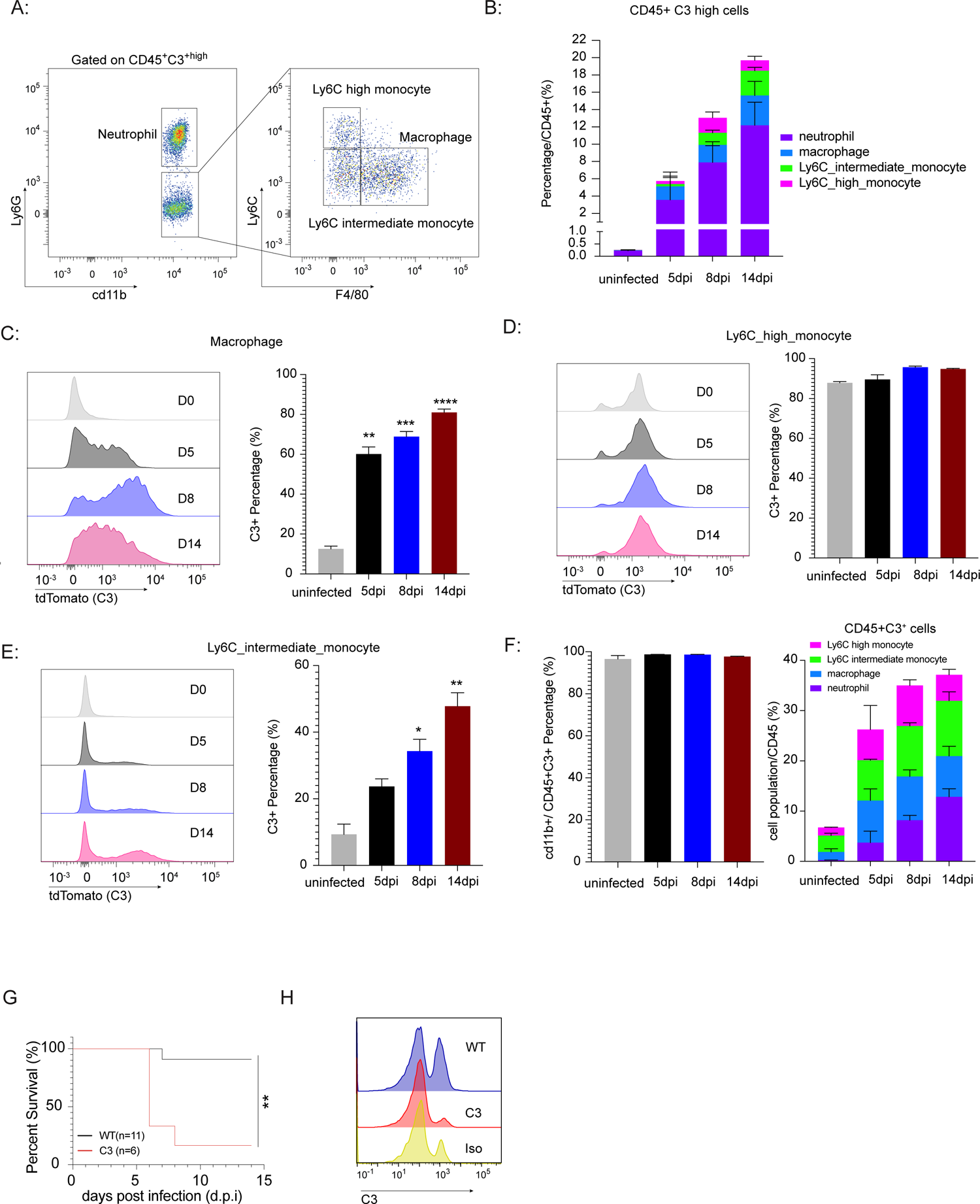
Luminal C3 is critical for the protection against EHEC and immune profiling of colonic CD45^+^C3^+^ cells during *C. rodentium* infection. **(A)** Flow cytometry gating of high-C3-expressing CD45^+^ colonic cells in the subepithelial compartment of C3^IRES-tdTomato^ reporter mice at day 14 post-*C. rodentium*-infection. (**B**). Percentage of high-C3-expressing neutrophils, macrophages, Ly6C intermediate- and Ly6C high-monocytes in CD45^+^ cells at days 0, 4, 5 and 14 post *Citrobacter rodentium* infection. **(C)** Representative histogram of C3 gene expression (MFI in C3-tdTomato reporter mice) in macrophages at days 0, 5, 8, and 14 post-infection in C3^IRES-tdTomato^ reporter mice, and percentages of C3-expressing macrophages in the total macrophage population from the colonic lamina propria at indicated days post-infection. **(D)** Representative histogram of C3 gene expression (MFI in C3-tdTomato reporter mice) in Ly6C-high monocytes at days 0, 5, 8, and 14 post-infection in C3^IRES-tdTomato^ reporter mice, and percentages of the C3-expressing Ly6C-high monocytes in the total Ly6C-high monocytes from the colonic lamina propria at indicated days post-infection. **(E)** Representative histogram of C3 gene expression (MFI in C3-tdTomato reporter mice) of Ly6C-intermediate monocytes at days 0, 5, 8, and 14 post-infection in C3^IRES-tdTomato^ reporter mice, and percentages of the C3-expressing Ly6C-intermediate monocytes in the total Ly6C-intermediate monocytes from colonic lamina propria at indicated days post-infection. **(F)** Percentages of CD11b^+^ cells amongst the total colonic lamina propria CD45^+^C3^+^ population at indicated days post-infection, and percentages of C3-expressing neutrophils, macrophages, Ly6C-intermediate and Ly6C-high monocytes in the CD45^+^ population at days 0, 5, 8 and 14 post-*Citrobacter rodentium*-infection. **(G)** Survival curve of C57BL/6 WT and C3-deficient mice (∼6-7 weeks old) following oral gavage with Enterohemorrhagic *E. coli* (EHEC) (10^9^ CFUs/mice). **(H)** Histogram of flow cytometric analysis of C3 opsonization of GFP^+^ EHEC at day 4 post-infection in WT or C3-deficient mice.

**Table S1:** Percentile abundances of species identified by group with ANCOM statistics.

**Table S2:** Spearman Correlation of bacterial genus with fecal C3 levels in healthy human gut microbiota.

**Table S3:** Sequencing primers and probes used in this study.

## Materials and Methods

### Mice

C57BL/6 mice were purchased from Jackson Laboratories, Taconic Biosciences, or Charles River Laboratories. B6. 129S4-C3^tm1Crr^/J, B6-C1qa^tm1d(EUCOMM)Wtsi/Tenn^J, B6.129S4-Mbl1^tm1Kata^ Mbl2^tm1Kata^/J mice were purchased from Jackson Laboratories. C3-tdTomato reporter (C3^IRES-tdTomato^ C57BL/6) mice were provided by Dr. Claudia Kemper (National Institute of Health). Mice were bred and housed in an SPF animal facility at Harvard Medical School (HMS). GF C57BL/6 mice were maintained at the Harvard Medical School Gnotobiotic Core Facility. Age-matched 6- to 12-week-old littermate male and female mice were used for experiments. All mice were used in accordance with animal care guidelines from Harvard Medical School Standing Committee on Animals and the National Institutes of Health.

### Antibiotic treatment and fecal transfer

Antibiotics were given via drinking water. For broad-spectrum antibiotic treatment (ABX), 0.5g/L of Vancomycin, 1g/L of Neomycin, 1g/L of Ampicillin and 1g/L of Metronidazole were added to drinking water. Antibiotic water was refreshed every three days. To confirm antibiotic activity, fecal matter was resuspended in PBS (1 ml/pellet) and cultured on Trypticase soy agar (TSA) and Brucella blood agar plates in aerobic and anaerobic conditions, respectively. For mouse microbiota fecal transfer, fecal samples collected from C57BL/6 WT mice from different vendors and resuspended in PBS at 100 mg/ml in the anaerobic chamber. Aliquots were kept at −80 °C. GF mice were orally gavaged with 200 μl of stock. For human microbiota fecal transfer, human fecal samples were collected and prepared as 25% glycerol stock as described (Alavi and Hsiao, STAR protocols, 2020). Fecal samples were normalized to be representative of 16S rRNA gene levels in 300ul of OD_600_=0.4 culture of defined community. GF mice were orally gavaged with 300ul normalized human fecal slurries.

### 16S rRNA gene sequencing of fecal samples and data analysis

Bacterial genomic DNA from frozen stool samples was extracted using phenol:chloroform:isoamyl alcohol and then purified further with the QIAquick PCR Purification kit (Qiagen)^45^. Purified DNA was quantified by Qubit dsDNA HS Assay (Thermo Fisher Cat# Q32854) and normalized. Amplicons were purified and quantified by Qubit dsDNA HS Assay and combined with equal mass to make a pooled library. The pooled library was multiplexed sequenced (Illumina MiSeq, 251 nt x 2 pair-end reads with 12 nt index reads) through Harvard’s Biopolymers Facility. Raw sequencing data was processed with QIIME2. In brief, raw sequencing data was imported to QIIME2 and demultiplexed, then DADA2 was used for sequence quality control and feature table construction. The feature table was used for beta diversity analysis, taxonomic analysis, and differential abundance testing using QIIME2. Beta group significance was determined by permutational analysis of variance (PERMANOVA). Identification of taxa associated with different groups was determined using Analysis of Composition of Microbiomes (ANCOM). Human fecal 16S rRNA gene library sequencing were described previously^46^. Spearman correlation between human gut microbiota at genus level and C3 levels was performed in R.

### Isolation of *Prevotella spp.* strains from Charles River (CR) mice gut microbiota

Fecal samples were collected from Charles River mice and homogenized in BBL thioglycollate media in an anaerobic chamber. Serial dilutions from the homogenized fecal sample were plated on Laked Brucella Blood Agar with Kanamycin and Vancomycin (LKV) Agar (Hardy Diagnostics) and cultured at 37°C in the anaerobic chamber for three days. Individual pigmented colonies were selected from these plates and cultured in peptone-yeast-glucose (PYG) broth for two days in the anaerobic chamber. Stocks were made from the liquid cultures and stored in −80°C. Bacterial gDNA were extracted from the liquid cultures using phenol:chloroform:isoamyl alcohol and then purified further with the QIAquick PCR Purification kit (Qiagen). Prevotella-specific 16S rRNA gene primers (gPrevo-F and R-)^14^ (Table S3) were used to confirm the stocks as *Prevotella spp.* and named as Prevotella CR1.

### C3 and C5 ELISA

C3 or C5 levels in feces or serum of mice were measured with a Mouse C3 ELISA kit (Abcam, ab157711) or a Mouse C5 ELISA kit (Abcam, ab264609) according to the manufacturer’s instructions. Briefly, fecal samples were collected and resuspended in dilution buffer at 100 mg/ml and centrifuged at 14000g for 10 minutes at 4°C and used the supernatant for C3/C5 levels measurement. Blood was collected from mice and centrifuged at 3000 rpm for 10 minutes after coagulation. Serum was collected for C3 concentration measurements.

### Gut permeability assay

A gut permeability assay was performed as previously described ^1–4^. Briefly, mice were fasted for four hours before oral gavage of 600 mg/kg FITC–3-5-kDa dextran (Sigma, FD4). Serum was collected six hours after gavage as well as control mice without gavage and analyzed using a Synergy HT plate reader (Bio-Tek) with 485 nm excitation and 530 nm emission.

### Preparation of intestinal cells and flow cytometry

Intestinal tissues were prepared as previously described.^47^ Briefly, distal colon and distal ileum were collected, treated with 30 mL of RPMI containing 1 mM dithiothreitol, 20 mM EDTA, and 2% FBS at 37°C for 15 min to isolate epithelial cells and intraepithelial lymphocytes. The intestinal tissues were then minced and dissociated in RPMI containing collagenase II (1.5 mg/mL; GIBCO), dispase (0.5 mg/mL), and 1% FBS, with constant stirring at 37°C (45 min for colon and small intestine). Single-cell suspensions were then filtered and washed with 10% FBS RPMI solution. Cells isolated above were stained with surface antibodies in a dilution of 1:200 on ice for 20 minutes in staining buffer (2% FBS in PBS) and washed twice with the same buffer. Cells were acquired with a BD FACSymphony (Beckman Coulter) or a BD LSR-II and analyzed in Flowjo.

### Stromal cells Isolation and culture

Colonic C3-expressing stromal cells were isolated from C3^IRES-tdTomato^ C57BL/6 mice. Briefly, single cell suspensions were prepared as described above from C3^IRES-tdTomato^ C57BL/6 mice. Live C3-expressing stromal cells were sorted as DAPI^-^Epcam^-^CD45^-^CD31^-^PDPN^+^tdTomato^+^ using a FACSAria (BD) into RPMI medium containing 10% FBS, then seeded on 96-well flat cell culture dishes (10,00 cells/well) in DMEM supplemented with 10% FBS and 1% penicillin/streptomycin at 37°C in the presence of 5% CO2. After 3 days of culture, live cells were collected, counted, and seeded as 10,000 cells/well in DMEM supplemented with 10% FBS and 1% penicillin/streptomycin at 37° overnight. Bacterial stimulants were fixed with 1% formalin for 24 hours and checked for no viability. Bacterial stimulants were normalized to OD_600_ 1 in RPMI medium and added 200ul to stromal cells, RPMI medium are used as control. LPS or MDP were dissolved in RPMI medium at the indicated concentrations and 200 ul were given to stromal cells.

### scRNASeq profiling of C3-expressing cells in colon

scRNASeq of colonic C3-expressing cells isolated from C3^IRES-tdTomato^ C57BL/6 mice was performed using 10x Genomics Chromium Next GEM Single Cell 3’ Reagent Kit (V3.1 chemistry). Briefly, colons from one male and one female C3^IRES-tdTomato^ C57BL/6 mice were collected, and single cell suspensions were prepared as described above. Live C3-expressing cells were sorted as DAPI^-^tdTomato^+^ using FACSAria (BD) into DMEM containing 10% FBS. A total of ∼10,000 C3-expressing cells from each mouse were collected and pooled together.

Samples were loaded into Chip G per the user guide from 10x Genomics. The Chip G was then run on a 10x Chromium Controller. 100ul of GEM emulsion was taken from the chip, and inspected visually for inconsistencies in the recovered volume, the uniformity of the emulsion, or the relative amount of partitioning oil, and incubated at RT in a thermocycler (45 min at 53°C, 5 min at 85°C, then held at 4°C until the next step). The emulsion was then broken by adding 125ul of Recovery Agent. The cDNA was then purified using Dynabeads MyOne SILANE. Per the 10x 3’v3.1 user guide, the 35ul of the purified cDNA was mixed with 15ul 10x cDNA primers (PN 200089) and 50ul of 10xAmp Mix (PN 2000047), then amplified on a thermocycler (98°C for 3 minutes, [98°C for 15 seconds, 63°C for 20 seconds, 72°C for 60 seconds], 12 repeats of the bracketed steps, 72°C for 60 seconds, then held at 4° C. The amplified cDNA was then purified and separated based on size using a 0.6x Beckman Coulter SPRIselect Reagent cleanup. The larger bound fragments in the bead pellet containing transcript-derived cDNA were eluted in 35ul of Qiagen Buffer EB for use in Gene Expression library construction. Library size was measured by an Agilent Bioanalyzer 2100 High Sensitivity DNA assay and quantified using a Qubit dsDNA HS Assay kit on a Qubit 4.0 Fluorometer.

A portion of the transcript-derived cDNA was fragmented, ligated to an Illumina Read 2 Sequencing Adaptor, and indexed with a unique 10x T Set A index, yielding the GEX library. 50ng of the transcript-derived cDNA was mixed with nuclease-free water up to a volume of 20ul, then mixed with 15ul of nuclease-free water, 5ul of 10x Fragmentation Buffer (PN 2000091), and 10ul of 10x Fragmentation Enzyme (PN 2000090), loaded onto a thermocycler, and incubated at 32°C for 2 minutes, then 65° C for 35 minutes, and then held at 4° C. The fragmented product was purified with a 0.6x SPRI, with 75ul of the supernatant saved. The supernatant was purified with a 0.8x SPRI, with the pellet-bound DNA saved and eluted in 50uL. All 50ul of eluted sample was mixed with 20ul 10x Ligation Buffer (PN 2000092), 10ul DNA ligase (PN 220110), and 20ul of Adaptor Oligos (2000094), then incubated on a thermocycler at 20C for 15min. This product was purified using a 0.8x SPRI cleanup, with the pellet-bound DNA saved and eluted in 30ul. These 30ul were then mixed with 20ul of a unique 10X T Set A index and 50ul of 10x Amp Mix, then incubated on a thermocycler at 98° C for 45 seconds, [98°C for 20 seconds, 54°C for 30 seconds, 72°C for 20 seconds], 13 repeats of the bracketed steps, 72°C for 60 seconds, then held at 4°C. The sample was then purified with a 0.6x SPRI, saving and transferring 150ul of the supernatant. The transferred supernatant was purified with a 0.8x SPRI and the pellet-bound DNA was eluted in 35ul. The finished library was measured on an Agilent BioAnalyzer and quantified on a Qubit 4.0. This library was then sequenced on an Illumina Novaseq SP 100, yielding approximately ∼366M reads in the GEX.

Raw sequencing data in the form of binary files were converted into FASTQ files using the mkfastq command from CellRanger v6.1.0. Then, the FASTQ files were demultiplexed into gene expression libraries using CellRanger’s count command, and only reads mapping unambiguously and with less than 2 mismatches were kept. For each single-cell library, reads were mapped to the mouse mm10 transcriptome using CellRanger. Duplicate reads, those mapping to multiple regions or having a low alignment score (MAPQ < 10) were filtered out. A final gene expression matrix with genes in rows and cells in columns was then constructed. Data were then analyzed using the Seurat R toolkit (https://satijalab.org/seurat/). Briefly, cells were first filtered based on the UMI count and mitochondrial gene percentage. Cells with < 500 UMIs or > 10 % mitochondrial genes were excluded from further analysis. Gene expression values for each cell were then normalized by the total expression, multiplied by an arbitrary scale factor of 10,000, and log-transformed. A principal component analysis (PCA) was performed on the top 200 most variable genes that were expressed in > 1% of the cells. The number of statistically significant PCs was determined by comparison with PCA over a randomized matrix as described previously^48^. The data were then visualized using the Uniform Manifold Approximation and Projection (UMAP) dimensionality reduction algorithm^49^ based on the significant PCs. Clustering was done on selected PCs using the FindNeighbors followed by the FindClusters functions, and the identity of the main clusters was determined based on the expression of common identifiers: Stromal (pdpn), epithelial (Epcam), myeloid (lyz2).

### Gene expression profiling by population RNAseq

A thousand live epithelial (EpCAM^+^), CD45^+^,and stromal cells (EpCAM^-^CD45^-^CD31^-^PDPN^+^) were doubly sorted from colons of WT SPF mice by BD FACS Aria II. Sorted cells were directly lysed with 5mL of TCL buffer (Qiagen, 1031576) containing 1% 2-mercaptoenthanol (Sigma-Aldrich, M6250), then flash frozen using dry-ice and stored at −80°C. Smart-Seq2 libraries for low-input RNA-seq were prepared by the Broad Technology Labs and were subsequently sequenced through the Broad Genomics Platform according to the Immgen protocol (https://www.immgen.org/img/Protocols/ImmGenULI_RNAseq_methods.pdf). The GCT and CLS files generated and used for downstream analysis. After normalization and filtered low count reads by DESeq2, fragments/counts per million were calculated for all the genes in the complement systems. Histogram and heatmaps with main components in complement system were depicted in Figure 3F, 6 G and H.

### Analysis of published single-cell data

For the dotplot gene expression profiles of complement system genes in human colonic cells, a full scRNASeq dataset of 428,000 intestinal cells^26^ was downloaded from https://www.gutcellatlas.org/, and then filtered as to keep cells with more than 500 genes, less than 25% mitochondrial reads, and genes with expression in more than three cells. A Scrublet (v.0.2.1) score cut-off of 0.25 was applied to exclude doublet. A global-scaling normalization method “LogNormalize” was applied to normalize the feature expression measurements for each cell by the total expression, multiplies this by a scale factor (10,000 by default), and log-transformed. Complement system gene expression table was extracted from the normalized table and plotted the expression levels using Dotplot in R.

### *Citrobacter rodentium* infection and C3-deposition analysis

*C. rodentium* DBS100 and GFP^+^ *C. rodentium* LB1 were provided by Dr. Lynn Bry (Brigham and Women’s Hospital).^30^ Survival studies were performed by orally infected 5×10^8^ CFU of *C. rodentium* DBS100 with twenty-one-day-old WT and C3^-/-^ C57BL/6 mice and followed for 28 days. Adult (8-12 weeks) WT and C3^-/-^ C57BL/6 mice were used to evaluate the roles of complement C3 in infection using *C. rodentium* doses between 5×10^8^-1X10^10^ CFU as indicated. Animal survival, weight, and disease symptoms were monitored daily for 3 weeks. Fecal burden represented by CFU/g of feces were monitored from day 7 to day 14. For C3 deposition analysis, age matched WT and C3^-/-^ mice were oral infected with 5×10^8^ CFU of GFP^+^ *C. rodentium* LB1. At day 7, fecal pellets were collected and resuspended in PBS containing a Roche proteinase inhibitor cocktail (Roche Cat. No. 04693159001). Cell suspensions were centrifuged at 50X g for 10 min at 4°C in order to pellet large particulate material. The supernatant fluids were filtered through a 40μm filter and the flow-through was then centrifuged to collect the bacterial cell pellet. Fecal bacteria were then ‘washed’ by resuspending the pellet in PBS, centrifuging, and resuspending with a proteinase inhibitor twice. The washed bacteria cells were resuspended in PBS containing 2% BSA and incubated for 30 minutes at 4°C before adding 1/10 volume of rabbit anti-mouse C3 antibody (sc-20137). After incubating the bacterial cells at 4^°^C for 1 hour, cells were washed twice and resuspended in PBS containing 1:10,000 diluted goat anti-rabbit IgG-647 antibody (Invitrogen). After 30mins at 4^°^C, bacterial cells were washed again and re-suspended in PBS before they were analyzed with a MACSQuant cytometer (Miltenyi Biotec). C3 deposition analysis in age-matched WT, MBL KO, C1qa KO mice were performed in similar procedure with DyLight405 (ab201798) conjugated recombinant rabbit monoclonal anti-mouse C3 antibody (ab259700).

### Cobra venom factor (CVF) treatment of wildtype mice

12.5ug CVF /25 gram body weight were given to WT mice via intraperitoneal injection one day before *C. rodentium* infection and on days 4, 8, 12 after infection. Control mice received saline injections. To test CVF depletion of C3, serum and fecal samples from CVF- or saline-treated mice were collected at day 0, day 1 and day 4 post CVF treatment to measure C3 levels using ELISA.

### Enterohemorrhagic *Escherichia Coli* (EHEC) infection

EHEC strain EDL931 were purchased from the American Type Culture Collection. The plasmid pUC18T-mini-Tn7T-Tp-gfpmut3^50^ was electroporated into EHEC EDL931 competent cells. The correct transformant was selected and confirmed to be positive for GFP by PCR as well as by flow cytometry, and designated as GFP^+^ EHEC. 6-7-week-old WT and C3-/-C57BL/6 mice were orally gavaged with Streptomycin (1mg/gram of body weight), then 1X10^9^ EHEC were given 24h after streptomycin treatment. Food was withdrawn 4 hours before giving EHEC and return soon after infection. For C3 deposition analysis, age-matched WT and C3-/-mice were orally infected with 1X10^9 CFU with GFP^+^ EHEC. At day 4 post-infection, fecal pellets were collected, and fecal bacteria were isolated as described earlier and stained with Dylight405 conjugated recombinant rabbit monoclonal anti-mouse C3 antibody (ab259700). GFP^+^ EHEC cells were as identified as FITC^+^ by flow cytometry on a MACSQuant cytometer.

### Mouse C3 RNAscope

Mouse C3 RNAScope *in situ* hybridization was performed using reagents and protocols from Advanced Cell Diagnostics. Briefly, colons from WT SPF mice were collected, fixed in 4% PFA, embedded in paraffin, and 5 μm sections were allowed to dry overnight. Sections were rehydrated twice in xylene for 5 min each, followed by two washes in 100% ethanol and one wash in 95% ethanol, 1 min each. After rehydration, the samples were incubated for 10 min in hydrogen peroxide, boiled in an antigen retrieval buffer for 15 min, followed by digestion with proteinase for 15 minutes at 40°C. Slides were washed twice with in situ hybridization (ISH) wash buffer, and hybridized with C3 probes (ACD#417841) for 2 hours at 40°C. After hybridization, amplification steps and staining were performed according to the protocol (RNAscope Fluorescent Reagent kit Assay).

### Immunofluorescent staining of C3-expressing cells in C3^IRES-tdTomato^ C57BL/6 mice

Distal colons were collected for staining. The intestines were opened and fixed in Silgard dish with 4% PFA at 4°C overnight. Then the tissues were dehydrated using 30% sucrose at 4°C overnight, then the tissues were embedded in OCT (Optimal cutting temperature compound) and frozen. 10μm sections were blocked with 10% donkey and goat serum in PBST (0.5% Triton-X100/PBS), then incubated with primary antibody (goat anti-tdTomato, 1:500; syrian hamster anti-pdpn, 1:500) overnight, followed by secondary antibody staining in PBST for 1 hour, and then stained with Hoechst for 5 mins, mounted with Prolong Antifade mounting medium, and sealed prior to imaging. Sections were imaged using a confocal microscopy at 63X and analyzed in Fiji. The C3-expressing cells isolated from C3^IRES-tdTomato^ C57BL/6 and WT mice were cultured and stained with goat anti-tdTomato in a 96-well plate with glass-like polymer bottom (Cellvis P96-1.5P) and imaged using a widefield microscopy to test anti-tdTomato antibody (**Figure S4A-C**).

### Bacterial probe staining

Colon tissues with fecal contents were collected and fixed by immersion in methacarn solution at 4°C for 48 hours. Tissues were then washed twice with 100% methanol (Sigma, MX0480-6), followed by twice with 100% ethanol (Fisher Scientific, BP2818500). Samples were then embedded in paraffin and cut into 4-μm-thick sections. In preparation for staining, the tissue paraffin slices were deparaffinized by heating at 60°C followed by immersion in four changes of room-temperature xylene (Sigma, 534056). The sample was then washed twice at room-temperature in 100% ethanol and rehydrated through a series of 95% then 70% ethanol incubations. The slices were then washed once with FISH-probe wash buffer^51^ for 5 minutes, and incubated with a probe mixture comprised of the *C. rodentium* rRNA FISH probe, the Eub338 probes, and FISH hybridization buffer (Moffitt et al, 2018). The samples were then incubated at 37 °C for 48 hours. Background fluorescence was removed by embedding the samples in a thin polyacrylamide gel and then cleared and treated with protease and detergent as described previously.^51^ Samples were then washed with 2× SSC five times for imaging immediately or stored at 4 °C.

### 16S-rRNA-specific probes for *C. rodentium*

A 30-nt *C. rodentium*-16S-rRNA-specific probe was designed using a previous pipeline^51^, and this sequence was concatenated with an additional 20-nt ‘readout’ sequence complementary to a previously designed fluorescently labeled FISH probe^51^. Two, 30-nt eubacterial FISH probes were designed by extending the standard Eub388 probe sequence to a total length of 30-nt, and concatenating different 20-nt ‘readout’ sequences. One Eub388 FISH probe carries a 5’ acrydite moiety, which served to anchor all eubacterial 16S rRNA during the tissue clearing process. As the sequence of the eubacterial probes do not overlap that of the *C. rodentium*-specific probe, we anticipated *C. rodentium* would be labeled by both.

### Bacterial imaging and data analysis

To prepare the sample for imaging, the sample was stained with two fluorescently-labeled readout probes complementary to the readout sequences on each of the *C. rodentium*-specific and Eub338 probes, as well as DAPI. The samples were imaged using buffers and protocols described previously^51^. Briefly, the samples were imaged on a home-built epi-fluorescence microscope using 95 mW of 750-nm illumination (Alexa750), 170mW of 635-nm illumination (Cy5), 26 mW of 545-nm illumination (orange fiducial beads), 77 mW of 473-nm illumination (Alexa488), and 72 mW of 408-nm illumination (DAPI) provided by a Celesta light engine (Lumencor). Illumination was reflected to the sample via a penta-band dichroic (Semrock FF421/491/567/659/776-Di01-25×36) and emitted light filtered with a penta-notch filter (Semrock FF01-391/477/549/639/741-25; FF01-441/511/593/684/817-25). Samples were imaged with a ×60 CFI PlanApo oil objective (Nikon) and two high-performance CMOS cameras (Hamamatsu, Flash 4.0). The mosaic for the cross-section colon slides was generated by stitching single field-of-views together. The epithelial boundaries were identified based on the 405-channel using Fiji. The *C. rodentium* signals were measured inside the epithelial boundaries by thresholding the *C. rodentium-*specific channel on intensity and summing the number of pixels above the threshold in Fiji. As different cross-sections had different sizes, the measured *C. rodentium* signals were normalized by the measured length of the epithelial-lumen boundary in each slice.

### Quantification And Statistical Analysis

Sample sizes for all experiments were chosen according to standard practice in the field. Statistical parameters including numbers, averages, deviation, and statistical tests are reported in the figures and corresponding legends. For all microscopy analysis, images were blinded prior to scoring. Data are represented as mean ± standard error (SEM) throughout the figures. Statistical analyses were performed in GraphPad Prism or R.

## Reference

1. Snyder, J.D., and Merson, M.H. (1982). The magnitude of the global problem of acute diarrhoeal disease: a review of active surveillance data. Bull World Health Organ 60, 605– 613.

2. Liu, L., Johnson, H.L., Cousens, S., Perin, J., Scott, S., Lawn, J.E., Rudan, I., Campbell, H., Cibulskis, R., Li, M., et al. (2012). Global, regional, and national causes of child mortality: an updated systematic analysis for 2010 with time trends since 2000. Lancet 379, 2151–2161. 10.1016/S0140-6736(12)60560-1.

3. Black, R., Fontaine, O., Lamberti, L., Bhan, M., Huicho, L., El Arifeen, S., Masanja, H., Walker, C.F., Mengestu, T.K., Pearson, L., et al. Drivers of the reduction in childhood diarrhea mortality 1980-2015 and interventions to eliminate preventable diarrhea deaths by 2030. J Glob Health 9. 10.7189/jogh.09.020801.

4. Guerrant, R.L., DeBoer, M.D., Moore, S.R., Scharf, R.J., and Lima, A.A.M. (2013). The impoverished gut—a triple burden of diarrhoea, stunting and chronic disease. Nat Rev Gastroenterol Hepatol 10, 220–229. 10.1038/nrgastro.2012.239.

5. Kunz, N., and Kemper, C. (2021). Complement Has Brains—Do Intracellular Complement and Immunometabolism Cooperate in Tissue Homeostasis and Behavior? Front. Immunol. 12. 10.3389/fimmu.2021.629986.

6. Noris, M., and Remuzzi, G. (2013). Overview of Complement Activation and Regulation. Seminars in Nephrology 33, 479–492. 10.1016/j.semnephrol.2013.08.001.

7. Yan, B., Freiwald, T., Chauss, D., Wang, L., West, E., Mirabelli, C., Zhang, C.J., Nichols, E.-M., Malik, N., Gregory, R., et al. (2021). SARS-CoV-2 drives JAK1/2-dependent local complement hyperactivation. Science Immunology 6. 10.1126/sciimmunol.abg0833.

8. Lloyd-Price, J., Arze, C., Ananthakrishnan, A.N., Schirmer, M., Avila-Pacheco, J., Poon, T.W., Andrews, E., Ajami, N.J., Bonham, K.S., Brislawn, C.J., et al. (2019). Multi-omics of the gut microbial ecosystem in inflammatory bowel diseases. Nature 569, 655–662. 10.1038/s41586-019-1237-9.

9. Ridaura, V.K., Faith, J.J., Rey, F.E., Cheng, J., Duncan, A.E., Kau, A.L., Griffin, N.W., Lombard, V., Henrissat, B., Bain, J.R., et al. (2013). Gut Microbiota from Twins Discordant for Obesity Modulate Metabolism in Mice. Science 341, 1241214. 10.1126/science.1241214.

10. Hsiao, E.Y., McBride, S.W., Hsien, S., Sharon, G., Hyde, E.R., McCue, T., Codelli, J.A., Chow, J., Reisman, S.E., and Petrosino, J.F. (2013). Microbiota modulate behavioral and physiological abnormalities associated with neurodevelopmental disorders. Cell 155, 1451– 1463.

11. Blanton, L.V., Charbonneau, M.R., Salih, T., Barratt, M.J., Venkatesh, S., Ilkaveya, O., Subramanian, S., Manary, M.J., Trehan, I., Jorgensen, J.M., et al. (2016). Gut bacteria that prevent growth impairments transmitted by microbiota from malnourished children. Science 351. 10.1126/science.aad3311.

12. Alavi, S., Mitchell, J.D., Cho, J.Y., Liu, R., Macbeth, J.C., and Hsiao, A. (2020). Interpersonal Gut Microbiome Variation Drives Susceptibility and Resistance to Cholera Infection. Cell 181, 1533–1546.e13. 10.1016/j.cell.2020.05.036.

13. Surana, N.K., and Kasper, D.L. (2017). Moving beyond microbiome-wide associations to causal microbe identification. Nature 552, 244–247. 10.1038/nature25019.

14. Gálvez, E.J.C., Iljazovic, A., Amend, L., Lesker, T.R., Renault, T., Thiemann, S., Hao, L., Roy, U., Gronow, A., Charpentier, E., et al. (2020). Distinct Polysaccharide Utilization Determines Interspecies Competition between Intestinal Prevotella spp. Cell Host & Microbe. 10.1016/j.chom.2020.09.012.

15. Alper, C.A., Johnson, A.M., Birtch, A.G., and Moore, F.D. (1969). Human C′3: Evidence for the Liver as the Primary Site of Synthesis. Science 163, 286–288. 10.1126/science.163.3864.286.

16. Morris, K.M., Aden, D.P., Knowles, B.B., and Colten, H.R. (1982). Complement biosynthesis by the human hepatoma-derived cell line HepG2. J Clin Invest 70, 906–913. 10.1172/jci110687.

17. Walport, M.J. (2001). Complement. N Engl J Med 344, 1058–1066. 10.1056/NEJM200104053441406.

18. Lubbers, R., van Essen, M.F., van Kooten, C., and Trouw, L.A. (2017). Production of complement components by cells of the immune system. Clin Exp Immunol 188, 183–194. 10.1111/cei.12952.

19. Liszewski, M.K., Kolev, M., Le Friec, G., Leung, M., Bertram, P.G., Fara, A.F., Subias, M., Pickering, M.C., Drouet, C., Meri, S., et al. (2013). Intracellular Complement Activation Sustains T Cell Homeostasis and Mediates Effector Differentiation. Immunity 39, 1143–1157. 10.1016/j.immuni.2013.10.018.

20. Pratt, J.R., Abe, K., Miyazaki, M., Zhou, W., and Sacks, S.H. (2000). In situ localization of C3 synthesis in experimental acute renal allograft rejection. Am J Pathol 157, 825–831. 10.1016/S0002-9440(10)64596-8.

21. Kulkarni, H.S., Elvington, M.L., Perng, Y.-C., Liszewski, M.K., Byers, D.E., Farkouh, C., Yusen, R.D., Lenschow, D.J., Brody, S.L., and Atkinson, J.P. (2019). Intracellular C3 Protects Human Airway Epithelial Cells from Stress-associated Cell Death. Am J Respir Cell Mol Biol 60, 144–157. 10.1165/rcmb.2017-0405OC.

22. Moon, R., Parikh, A.A., Szabo, C., Fischer, J.E., Salzman, A.L., and Hasselgren, P.O. (1997). Complement C3 production in human intestinal epithelial cells is regulated by interleukin 1beta and tumor necrosis factor alpha. Arch Surg 132, 1289–1293. 10.1001/archsurg.1997.01430360035007.

23. Andoh, A., Fujiyama, Y., Bamba, T., and Hosoda, S. (1993). Differential cytokine regulation of complement C3, C4, and factor B synthesis in human intestinal epithelial cell line, Caco-2. J Immunol 151, 4239–4247.

24. Sünderhauf, A., Skibbe, K., Preisker, S., Ebbert, K., Verschoor, A., Karsten, C.M., Kemper, C., Huber-Lang, M., Basic, M., Bleich, A., et al. (2017). Regulation of epithelial cell expressed C3 in the intestine – Relevance for the pathophysiology of inflammatory bowel disease? Molecular Immunology 90, 227–238. 10.1016/j.molimm.2017.08.003.

25. Kolev, M., West, E.E., Kunz, N., Chauss, D., Moseman, E.A., Rahman, J., Freiwald, T., Balmer, M.L., Lötscher, J., Dimeloe, S., et al. (2020). Diapedesis-Induced Integrin Signaling via LFA-1 Facilitates Tissue Immunity by Inducing Intrinsic Complement C3 Expression in Immune Cells. Immunity 52, 513–527.e8. 10.1016/j.immuni.2020.02.006.

26. Elmentaite, R., Kumasaka, N., Roberts, K., Fleming, A., Dann, E., King, H.W., Kleshchevnikov, V., Dabrowska, M., Pritchard, S., Bolt, L., et al. (2021). Cells of the human intestinal tract mapped across space and time. Nature 597, 250–255. 10.1038/s41586-021-03852-1.

27. Pendse, M., Li, Y., Salinas, C.N., Quinn, G., Vo, N., Propheter, D.C., Dende, C., Crofts, A.A., Koo, E., Hassell, B., et al. (2022). Macrophages regulate gastrointestinal motility through complement component 1q (Immunology) 10.1101/2022.01.27.478097.

28. Bouskra, D., Brézillon, C., Bérard, M., Werts, C., Varona, R., Boneca, I.G., and Eberl, G. (2008). Lymphoid tissue genesis induced by commensals through NOD1 regulates intestinal homeostasis. Nature 456, 507–510. 10.1038/nature07450.

29. Mullineaux-Sanders, C., Sanchez-Garrido, J., Hopkins, E.G.D., Shenoy, A.R., Barry, R., and Frankel, G. (2019). Citrobacter rodentium–host–microbiota interactions: immunity, bioenergetics and metabolism. Nat Rev Microbiol 17, 701–715. 10.1038/s41579-019-0252-z.

30. Belzer, C., Liu, Q., Carroll, M.C., and Bry, L. (2011). The role of specific IgG and complement in combating a primary mucosal infection of the gut epithelium. Eur J Microbiol Immunol (Bp) 1, 311–318. 10.1556/EuJMI.1.2011.4.7.

31. Silberger, D.J., Zindl, C.L., and Weaver, C.T. (2017). Citrobacter rodentium: A Model Enteropathogen for Understanding the Interplay of Innate and Adaptive Components of Type 3 Immunity. Mucosal Immunol 10, 1108–1117. 10.1038/mi.2017.47.

32. Deng, W., Li, Y., Vallance, B.A., and Finlay, B.B. (2001). Locus of Enterocyte Effacement from Citrobacter rodentium: Sequence Analysis and Evidence for Horizontal Transfer among Attaching and Effacing Pathogens. Infect Immun 69, 6323–6335. 10.1128/IAI.69.10.6323-6335.2001.

33. Caulfield, L.E., de Onis, M., Blössner, M., and Black, R.E. (2004). Undernutrition as an underlying cause of child deaths associated with diarrhea, pneumonia, malaria, and measles. The American Journal of Clinical Nutrition 80, 193–198. 10.1093/ajcn/80.1.193.

34. Chen, R.Y., Mostafa, I., Hibberd, M.C., Das, S., Mahfuz, M., Naila, N.N., Islam, M.M., Huq, S., Alam, M.A., Zaman, M.U., et al. (2021). A Microbiota-Directed Food Intervention for Undernourished Children. N Engl J Med 384, 1517–1528. 10.1056/NEJMoa2023294.

35. Charbonneau, M.R., O’Donnell, D., Blanton, L.V., Totten, S.M., Davis, J.C.C., Barratt, M.J., Cheng, J., Guruge, J., Talcott, M., Bain, J.R., et al. (2016). Sialylated Milk Oligosaccharides Promote Microbiota-Dependent Growth in Models of Infant Undernutrition. Cell 164, 859– 871. 10.1016/j.cell.2016.01.024.

36. Halstensen, T.S., and Brandtzaeg, P. (1991). Local complement activation in inflammatory bowel disease. Immunol Res 10, 485–492. 10.1007/BF02919746.

37. Krausgruber, T., Fortelny, N., Fife-Gernedl, V., Senekowitsch, M., Schuster, L.C., Lercher, A., Nemc, A., Schmidl, C., Rendeiro, A.F., Bergthaler, A., et al. (2020). Structural cells are key regulators of organ-specific immune responses. Nature 583, 296–302. 10.1038/s41586-020-2424-4.

38. Davidson, S., Coles, M., Thomas, T., Kollias, G., Ludewig, B., Turley, S., Brenner, M., and Buckley, C.D. (2021). Fibroblasts as immune regulators in infection, inflammation and cancer. Nat Rev Immunol 21, 704–717. 10.1038/s41577-021-00540-z.

39. Costa, A., Kieffer, Y., Scholer-Dahirel, A., Pelon, F., Bourachot, B., Cardon, M., Sirven, P., Magagna, I., Fuhrmann, L., Bernard, C., et al. (2018). Fibroblast Heterogeneity and Immunosuppressive Environment in Human Breast Cancer. Cancer Cell 33, 463–479.e10. 10.1016/j.ccell.2018.01.011.

40. Elyada, E., Bolisetty, M., Laise, P., Flynn, W.F., Courtois, E.T., Burkhart, R.A., Teinor, J.A., Belleau, P., Biffi, G., Lucito, M.S., et al. (2019). Cross-Species Single-Cell Analysis of Pancreatic Ductal Adenocarcinoma Reveals Antigen-Presenting Cancer-Associated Fibroblasts. Cancer Discovery 9, 1102–1123. 10.1158/2159-8290.CD-19-0094.

41. Puram, S.V., Tirosh, I., Parikh, A.S., Patel, A.P., Yizhak, K., Gillespie, S., Rodman, C., Luo, C.L., Mroz, E.A., Emerick, K.S., et al. (2017). Single-Cell Transcriptomic Analysis of Primary and Metastatic Tumor Ecosystems in Head and Neck Cancer. Cell 171, 1611–1624.e24. 10.1016/j.cell.2017.10.044.

42. Tirosh, I., Izar, B., Prakadan, S.M., Wadsworth, M.H., Treacy, D., Trombetta, J.J., Rotem, A., Rodman, C., Lian, C., Murphy, G., et al. (2016). Dissecting the multicellular ecosystem of metastatic melanoma by single-cell RNA-seq. Science 352, 189–196. 10.1126/science.aad0501.

43. Calon, A., Lonardo, E., Berenguer-Llergo, A., Espinet, E., Hernando-Momblona, X., Iglesias, M., Sevillano, M., Palomo-Ponce, S., Tauriello, D.V.F., Byrom, D., et al. (2015). Stromal gene expression defines poor-prognosis subtypes in colorectal cancer. Nat Genet 47, 320–329. 10.1038/ng.3225.

44. Royer, D.J., Echegaray-Mendez, J., Lin, L., Gmyrek, G.B., Mathew, R., Saban, D.R., Perez, V.L., and Carr, D.J. Complement and CD4+ T cells drive context-specific corneal sensory neuropathy. eLife 8, e48378. 10.7554/eLife.48378.

45. Wu, M., McNulty, N.P., Rodionov, D.A., Khoroshkin, M.S., Griffin, N.W., Cheng, J., Latreille, P., Kerstetter, R.A., Terrapon, N., Henrissat, B., et al. (2015). Genetic determinants of in vivo fitness and diet responsiveness in multiple human gut Bacteroides. Science 350, aac5992. 10.1126/science.aac5992.

46. Alavi, S., and Hsiao, A. (2020). Protocol for Microbiome Transplantation in Suckling Mice during Vibrio cholerae Infection to Study Commensal-Pathogen Interactions. STAR Protocols 1, 100200. 10.1016/j.xpro.2020.100200.

47. Geva-Zatorsky, N., Sefik, E., Kua, L., Pasman, L., Tan, T.G., Ortiz-Lopez, A., Yanortsang, T.B., Yang, L., Jupp, R., Mathis, D., et al. (2017). Mining the Human Gut Microbiota for Immunomodulatory Organisms. Cell 168, 928–943.e11. 10.1016/j.cell.2017.01.022.

48. Klein, A.M., Mazutis, L., Akartuna, I., Tallapragada, N., Veres, A., Li, V., Peshkin, L., Weitz, D.A., and Kirschner, M.W. (2015). Droplet Barcoding for Single-Cell Transcriptomics Applied to Embryonic Stem Cells. Cell 161, 1187–1201. 10.1016/j.cell.2015.04.044.

49. McInnes, L., Healy, J., and Melville, J. (2020). UMAP: Uniform Manifold Approximation and Projection for Dimension Reduction. 10.48550/arXiv.1802.03426.

50. Choi, K.-H., and Schweizer, H.P. (2006). mini-Tn7 insertion in bacteria with single attTn7 sites: example Pseudomonas aeruginosa. Nat Protoc 1, 153–161. 10.1038/nprot.2006.24.

51. Moffitt, J.R., Bambah-Mukku, D., Eichhorn, S.W., Vaughn, E., Shekhar, K., Perez, J.D., Rubinstein, N.D., Hao, J., Regev, A., Dulac, C., et al. (2018). Molecular, spatial, and functional single-cell profiling of the hypothalamic preoptic region. Science 362, eaau5324. 10.1126/science.aau5324.

